# Silicon exaptation discovers antimicrobial peptides

**DOI:** 10.64898/2026.09.21.753171

**Authors:** Gabriela Wojtan, Noora Ahonen, Burcu Firatligil Yildirir, Sofia Julin, Mirabela Babin, Timo Laakko, Christian Ganser, Feng-Yueh Chan, Arola Suvi, Nonappa, Christopher Jonkergouw, Ali Miserez, Takayuki Uchihashi, Piotr Batys, Pezhman Mohammadi

**Author notes:** Authors contributed equally (Gabriela Wojtan, Noora Ahonen).

## Abstract

Advances in machine learning (ML) have enabled the *de novo* design of synthetic proteins and peptides tailored for specific structural or functional properties. As ML becomes a central tool in forward design, we suggest that its value extends beyond optimizing known objectives. It can also lead to what we term here in this work as “silicon exaptation”, revealing unexpected functional properties that were never part of the original design goals. This reflects a principle common in biology, that repurposes traits where features evolved for one role can later enable unforeseen functions. In this context, these accidental findings are not constrained by preconceptions but arise from latent function emergence and the algorithm’s gradual drift, a new process that can co-evolves with human curiosity, intuition, and interpretive reasoning, which can be used to recognize unexpected patterns and transform them into new discoveries. In this study, we discovered a set of peptides through silicon exaptation, in which the ML pipeline was originally optimized for structural characteristics unrelated to any antimicrobial functionalities. The peptides were originally designed to adopt defined secondary structures in response to environmental stimuli, with no antimicrobial properties intended or included in the training objectives. We systematically evaluated the antimicrobial potential of these peptides using broth microdilution assays and membrane integrity tests, identifying several candidates with potent and selective antibacterial activity without detectable cytotoxicity toward mammalian cells.

## Introduction

The integration of machine learning (ML) into biomolecular research has transformed the way peptides and proteins are engineered for various medical and industrial applications.^1–6^ By learning sequence–structure–function relationships from large experimental or simulated datasets, ML models now enable *de novo* prediction and forward design of sequences tailored for targeted properties. ^7–9^ Recent applications span structural stabilization, folding prediction, enzyme design, and therapeutic discovery. Importantly, these data-driven approaches differ from traditional trial-and-error strategies in their ability to generalize across high-dimensional sequence space and generate viable candidates with reduced experimental burden, costs and time.^10–12^

Within peptide science, the dominant design paradigm has historically been application driven. Antimicrobial peptides (AMPs) are good examples.^13,14^ The urgency for the use of AMPs has been amplified by the global crisis of multidrug-resistant bacteria, which are becoming substantially difficult to treat and have created an acute demand for alternative therapeutic strategies in recent years.^15–17^ Over the past decades, intensive research has sought to identify natural sequence motifs, their physicochemical parameters, and key structural determinants that promote efficient bacterial membrane disruption, complemented by rational engineering of AMPs to enhance their activity.^18–21^ In recent years, ML models trained on protein/peptide sequence-structural databases and curated AMP databases have refined these objectives, yielding algorithms that can classify and identify new previously undiscovered AMPs.^22–24^ Consequently, the future trend of AMP design workflows is likely to be increasingly integrated with advanced ML and computational modeling. This enables accelerated optimization toward predefined functional endpoints such as cationicity, hydrophobicity, amphipathicity, and other properties explicitly correlate with antimicrobial potency with a specific goal of discovering and engineering novel *de novo* classes of AMPs.^25,26^

Despite the high success rate of such approaches, objective-driven design explores only a narrow, low-entropy corridor of peptide sequence space defined by known AMP descriptors. Yet peptide function resides on a richer latent manifold where structural constraints, environmental responsiveness, and mesoscale assembly couple nonlinearly to activity. This raises two key questions: a) can navigating these broader manifolds uncover antimicrobial activity as an emergent property rather than a targeted design outcome? and b) if so, does this imply that ML design should be re-envisioned not as a statistical optimization tool, but also as an instrument for discovering unexpected functional traits?

In this regard nature provides compelling inspiration. From an evolutionary perspective, biology routinely repurposes traits through a process known as exaptation, in which features that evolved for one role later enable one or multiple different unforeseen functions.^27^ By analogy, we propose that ML pipelines can recapitulate this logic through a similar strategy we term “Silicon Exaptation”, revealing unexpected functional properties that were never part of the original design goals leading to ML-enabled serendipitous discovery. For example, sequence patterns that stabilize conformation, modulate amphipathicity in response to environmental changes, or tune interfacial binding may incidentally meet the biophysical requirements for bacterial membrane disruption. In this regime, antimicrobial activity can appear as a latent byproduct of sequence patterns optimized for other goals (e.g., structural control, stimulus responsiveness, higher order supramolecular self-assemblies). By traversing off-objective neighborhoods, ML-guided search can therefore enable discovery of new candidates.

Motivated by this fresh perspective, we set out to test whether the ML pipeline trained for structural objectives from our earlier work could surface such hidden functions (**Fig. 1**). To test this, we used a framework designed to explore sequence features favoring defined secondary structures under specific solution conditions, with no initial antimicrobial objectives. The initial rationale was to expand beta-hairpins design space for conformational control and environment-responsive folding, providing a versatile platform for material science rather than therapeutic applications in our previous work (**Fig .1**). While the peptides were designed exclusively for structural responsiveness, unexpectedly, several of the designed peptides displayed antimicrobial activity, despite no such objective being encoded demonstrating the unexpected potential for function emergence in ML-driven peptide engineering. We therefore subjected these peptides to a comprehensive experimental and computational characterizations, combining antimicrobial susceptibility assays, membrane integrity and cytotoxicity measurements, *in situ* nanoscale imaging of membrane remodeling, and molecular dynamics simulations of peptide–membrane interactions. Unexpectedly, this analysis revealed a highly selective antimicrobial mechanism directed toward bacterial membranes, while sparing mammalian cells, and uncovered a distinct sequence of events involving surface saturation, membrane-targeted rupture, and rapid population collapse. Together, these findings led us to propose a hybrid Throttle–Order–Rupture–Collapse (TORC) mechanism, providing a mechanistic framework for understanding the emergent antimicrobial activity of these peptides.

**Figure 1.**
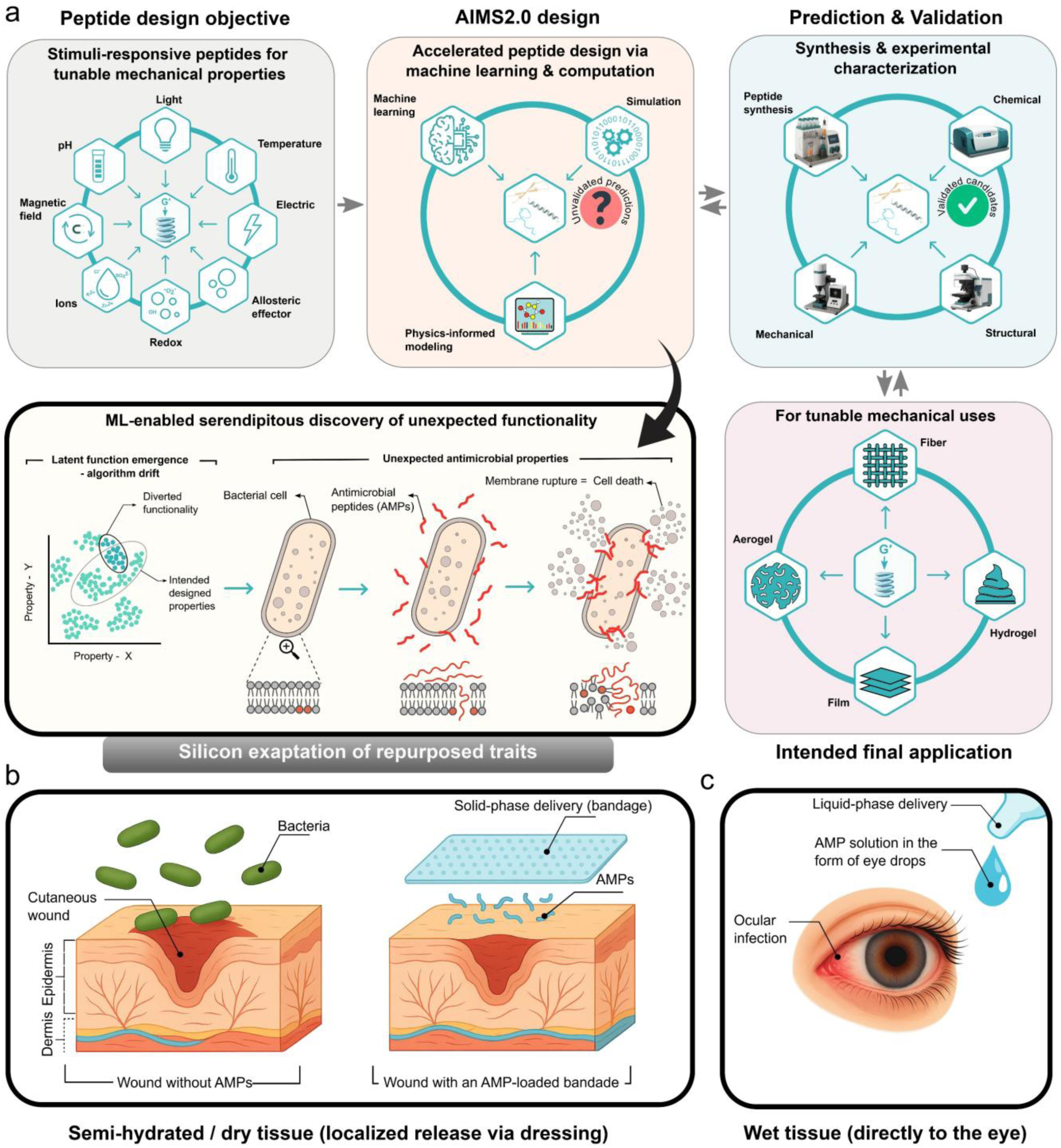
Schematic representation of Silicon exaptation of repurposed traits in ML-enabled serendipitous discovery of antimicrobial activity. **a)** Workflow of peptide design objectives and applications. Stimuli-responsive peptides originally designed for tunable mechanical properties. Accelerated peptide design performed with AIM2.0, integrating machine learning, physics-informed modeling, and simulations. Peptides were synthesized and experimentally validated through chemical, structural, and mechanical characterization resulting in peptides with potential use for various material formats such as fibers, aerogels, films, and hydrogels. Unexpectedly, ML-driven design led to the emergence of latent functions not included in the original objectives, revealing antimicrobial peptides (AMPs) with activity against bacterial membranes. **b and c)** Discovered AMPs retained antimicrobial activity in both semi-hydrated/dry tissue environments and fully wet tissue environments. This represents promising candidates for broad biomedical applications across different physiological conditions such as cutaneous wound treatment with AMP-loaded dressings, panel b, or in ocular infection treatment with AMP-containing eye drops, panel c.

## Results and discussion

### Original purpose of designer peptides

The peptide library investigated here originated from our efforts to design β-hairpin scaffolds with tunable conformational switching for material science applications accelerated through ML and computational modeling (**Fig.1 and table S1**).^28^ Sequences were constructed according to β-hairpin engineering principles by alternating hydrophobic and cationic residues to drive strand amphiphilicity, turn-inducing dipeptides (e.g., DK, GD) to nucleate β-turns, and side-chain patterning to stabilize strand registry and folding. Their original design was guided by biomimetic principles of β-hairpin folding inspired by high performance structural proteins such as fibroin and resilin. The main goal was engineering sequences capable of undergoing controlled reversible conformational switching (disorder-to-β-sheet conversion) in response to an external small molecular inducer such as SDS, thereby creating adaptive building blocks for supramolecular materials with tunable mechanical properties rather than therapeutic activity. To facilitate systematic exploration, each peptide was assigned a structured identifier. The prefix AI.PBH denotes Artificial Intelligence–designed Peptide Beta-Hairpin. This is followed by a number indicating the peptide length (e.g., 19, 25, 33, 39) and then by a unique identifier generated by the AIMS 2.0 pipeline (e.g., 016, 030). For example, AI.PBH_25_016 refers to a 25– amino acid β-hairpin peptide, the 016 corresponds to unique sequence generated in that design batch reflecting on the design parameters and computational origin.

### Growth inhibition and unexpected antimicrobial activity

We experimentally tested all 49 peptides against gram-negative bacteria in broth microdilution assays and monitored bacterial growth kinetics over 16 hours (**Fig. 2**). Growth was measured using optical density measurement at 600 nm (OD600). Thirteen peptides completely suppress the bacterial proliferation and OD increase, indicating potent growth inhibition under these conditions as evidenced by flat or near-baseline OD600 curves comparable to the positive control. Some of these peptides with clear inhibitory effects include AI.PBH_25_016, AI.PBH_25_114, AI.PBH_39_006, and AI.PBH_25_109. In other cases (e.g., AI.PBH_33_081, AI.PBH_19_025, AI.PBH_39_004), growth was delayed or attenuated, suggesting partial bacteriostatic activity. In contrast, many sequences—including AI.PBH_19_059, AI.PBH_25_018, AI.PBH_37_034, and AI.PBH_39_018 displayed growth kinetics indistinguishable from untreated control, indicating lack of any measurable activity at the tested concentrations.

**Figure 2.**
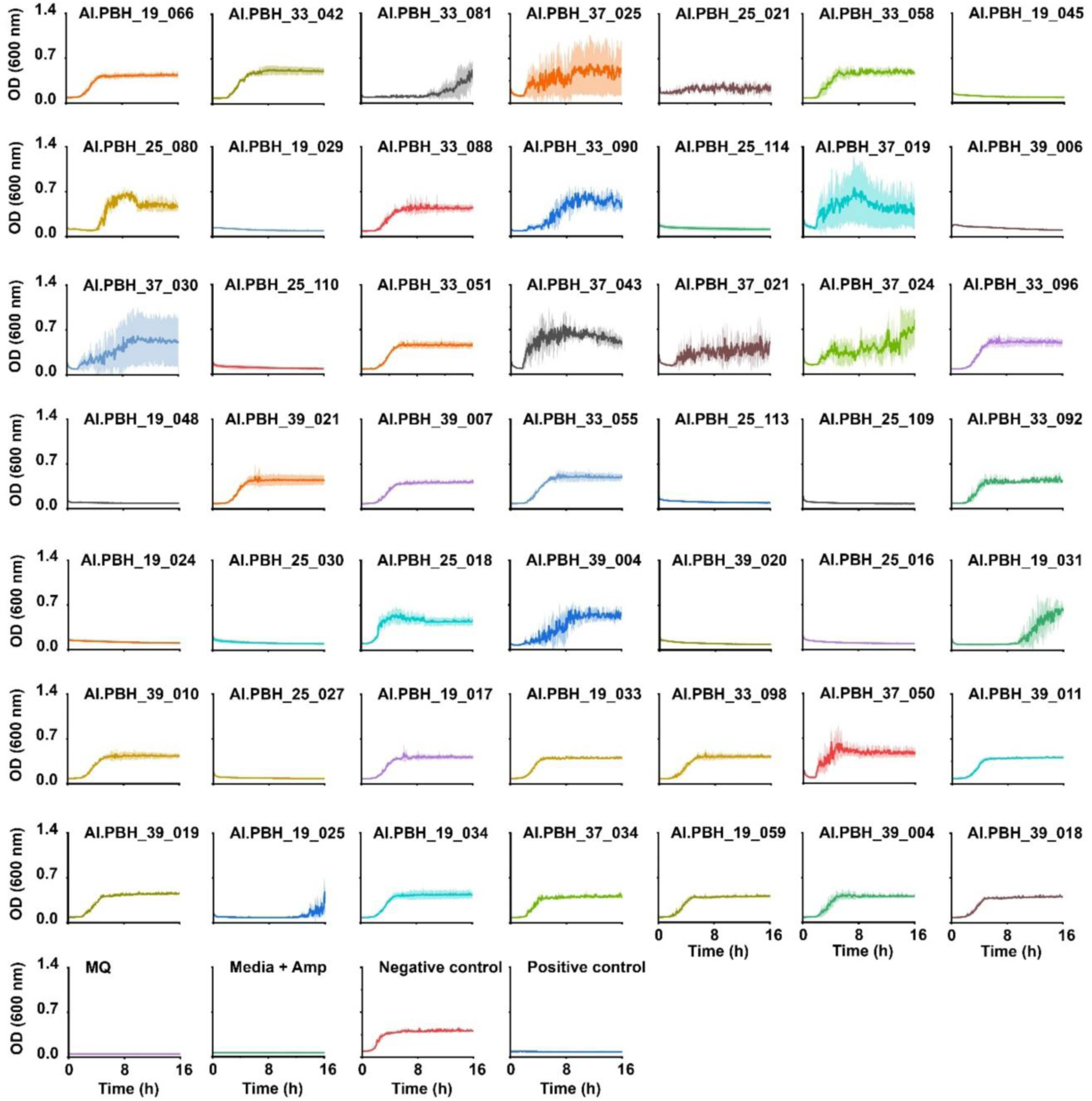
Emergent antimicrobial activity of 43 ML-designed peptides. The panels show growth curves (OD₆₀₀) for the period of 1c hours for all forty-nine. Each panel shows the result for one peptide (ID in title) at 350 µM; the bottom row shows controls: Milli-Q water (blank), medium + ampicillin only, negative control (bacteria + vehicle), and positive control (bacteria + ampicillin). Solid lines represent the mean, and shading represents the standard deviation. Measurement for inactive peptides repeated three times (N=3), and for active peptides six times (N=c).

### Static antimicrobial activity from agar diffusion

Building on the 16-hour broth microdilution kinetics (**Fig. 2**), we next quantified static antimicrobial activity under diffusion-limited, non-agitated conditions using agar diffusion. Representative plates (**Fig. 3a**) show well-defined zones of inhibition surrounding 13 active peptides. Zones of inhibition diameters were calculated from calibrated plate images and summarized for each active peptide across replicates (**Fig. 3b**), revealing robust potency for all peptides with minor variability. The ranking activity did not directly follow the trend in broth kinetics, supporting differences in intrinsic activity; minor deviations are consistent with assay-specific effects, such as differential diffusivity, peptide–agar interactions, and sensitivity to medium composition (e.g., ionic strength, divalent cations, agar), that can attenuate halo size independently of bacteriostatic potency. Taken together, the plate-based readout confirms peptide inhibitory activity under static conditions and complements the broth measurements.

**Figure 3.**
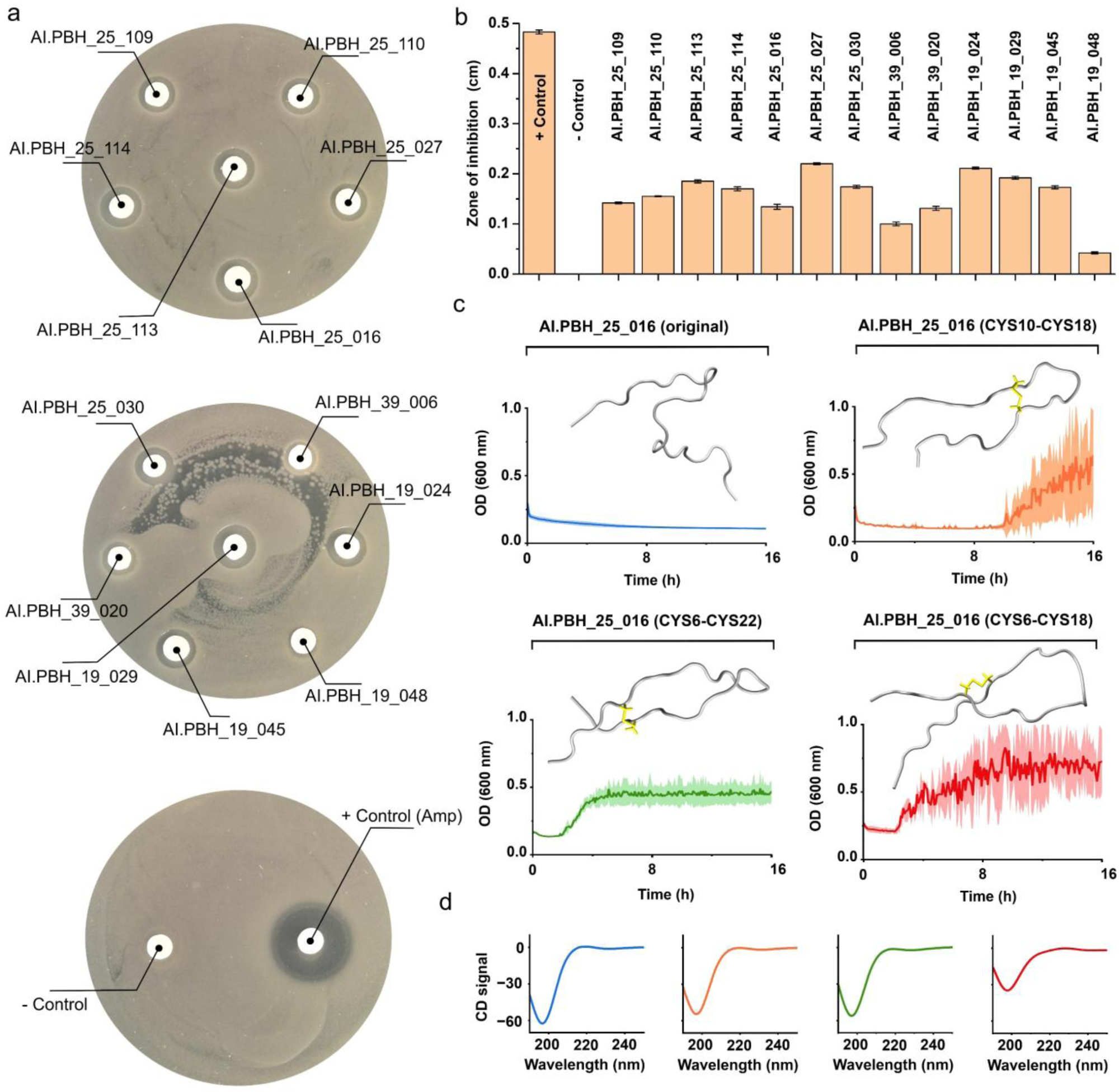
Static activity and structure–function trade-off reducing peptide mobility decreases inhibition potency. **a)** Representative agar disc-diffusion assay and quantification. Sterile paper discs loaded with individual peptides (peptide IDs labeled). Ampicillin was used as a positive control (+Control, Amp); empty discs were the negative control (–Control). **b**) After the tests, zones of inhibition were measured; the bar plot shows inhibition diameters (cm) for each peptide and controls (N=4). **c**) Growth inhibition assays of E. coli in liquid culture with peptide AI.PBH_25_01c and its cysteine mutants carrying intramolecular disulfide bonds (yellow). The panel shows representative structural schematics (top) illustrate the position of engineered disulfide bonds (yellow) and corresponding bacterial growth curves (bottom) monitored at ODc00 over 1c h. All measurements are carried out three times (N=3). **d**) The corresponding CD measurements for each peptide as in c panel. From left to right AI.PBH_25_01c (original), AI.PBH_25_01c (CYS10–CYS18), AI.PBH_25_01c (CYSc–CYS22) and AI.PBH_25_01c (CYSc–CYS18).

### Mobility-enabled activity from β-hairpin–designed peptides

Guided by our silicon exaptation perspective, we asked whether the unexpected antimicrobial activity uncovered by our structure-focused ML pipeline depends on conformational mobility rather than a predefined fold. Although all the peptides in this study were originally designed to adopt a β-hairpin under specific solution cues, our measurements indicate this is not the case under the assay conditions. Because AI.PBH_25_016 was the top performer in our screen, we used it as a mechanistic probe. In liquid culture, the unconstrained AI.PBH_25_016 free of intramolecular restraints produced rapid and sustained growth inhibition (**Fig. 3c, Fig. S1**). In contrast, variants engineered to restrict backbone motion via intramolecular Cys–Cys constraints (CYS10–CYS18, CYS6–CYS22, CYS6–CYS18) substitutions showed markedly reduced inhibition, with cultures recovering over time. Thus, limiting conformational freedom consistently diminishes the potency.

To determine whether the activity loss reflects stabilization of a particular secondary structure, we collected far-UV circular dichroism (CD) spectra for all constructs (**Fig. 3d**). Contrary to the β-hairpin design intent, none of the variants displayed a β-sheet/hairpin signature (strong negative band at ∼215–218 nm with a compensating positive at ∼195 nm). Instead, all peptides exhibited intrinsically disordered profiles, dominated by a single negative band near ∼200 nm. Together, these results support a model in which dynamic, disordered ensembles enable adaptive, multivalent engagement with gram-negative membranes, while covalent or electrostatic constraints narrow the accessible conformational space and limit essential adaptive interactions.

### Sequence features distinguishing active and inactive *de novo* peptides

We then investigated sequence design for 13 active and 36 inactive peptides (**Fig. 2 and Table S1**). Despite this evident functional divergence, both groups shared several clear similarities. All peptides were highly soluble in water and physiological buffers, excluding solubility as a discriminating factor. In addition, both active and inactive sequences were strongly enriched in lysine, consistent with the canonical cationic profile of antimicrobial peptides (**Fig. 4a**). On average, lysine accounted for ∼45–55% of residues in active sequences and ∼35–45% in inactive ones, indicating that while lysine-richness is necessary, it is not alone sufficient for antimicrobial function. A more detailed compositional analysis revealed critical differences. Active peptides consistently maintained a high net positive charge, typically with a basic-to-acidic residue ratio of > 3:1 (**Fig. 4a and Table S1**). In contrast, inactive sequences exhibited a more balanced ratio, often closer to 2:1 or lower, reflecting a greater prevalence of negatively charged residues (aspartic acid and glutamic acid). Quantitatively, the active group contained on average about 8–10% acidic residues, whereas the inactive group had nearly double that proportion (15–20%). This enrichment of acidic residues in the inactive group likely reduces net cationicity and weakens electrostatic interactions with negatively charged bacterial membranes.

**Figure 4.**
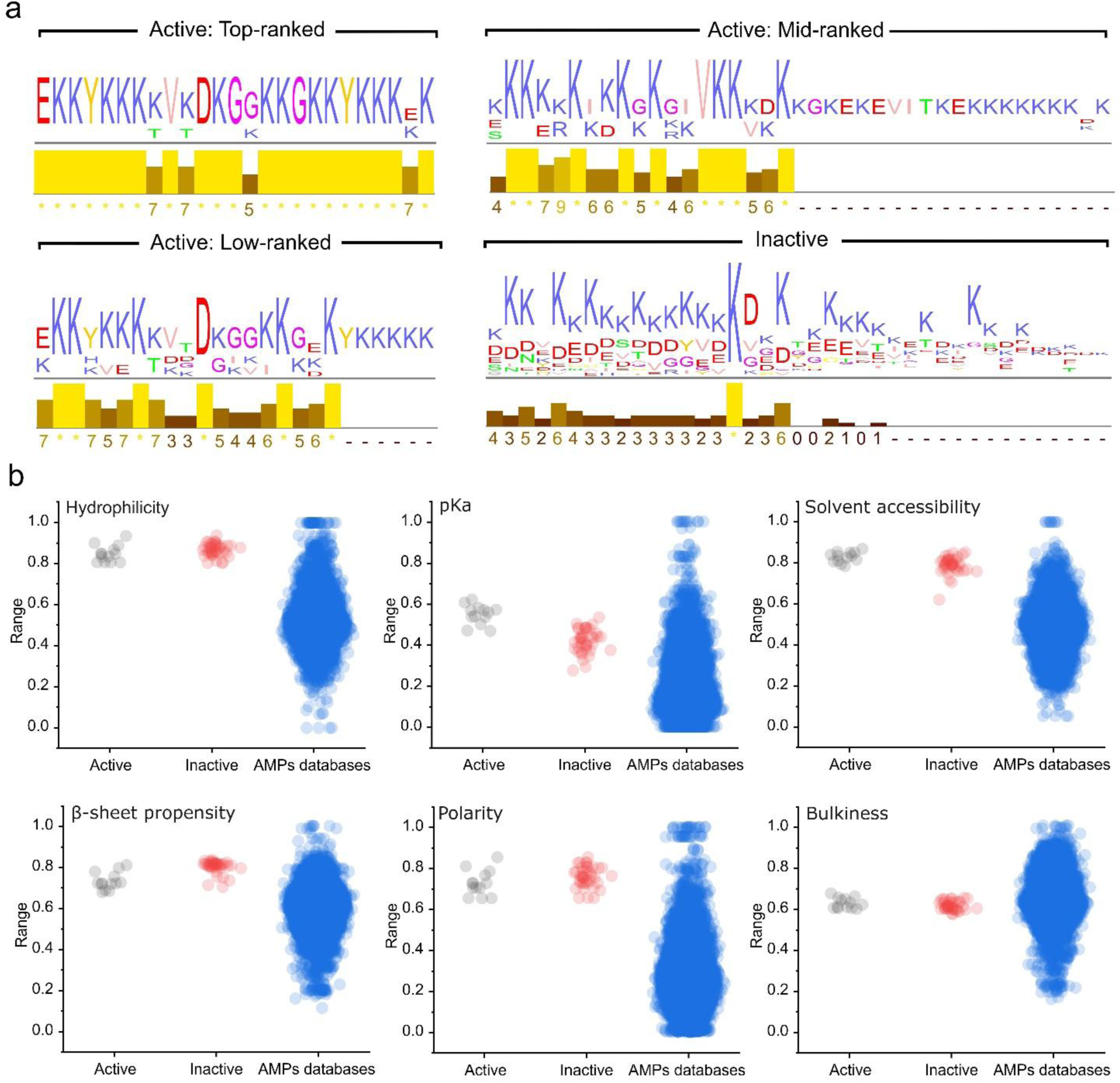
Comparative physicochemical and structural properties of peptides derived from in silico exaptation and trait repurposing, alongside 81,424 peptides from AMP databases. **a)** Sequence logos ranked by activity. **b)** Panels show distributions of six peptide descriptors including hydrophilicity, pKa, solvent accessibility, β-sheet propensity, polarity, and bulkiness for peptide with active antimicrobial property (Active, grey), without detectable activity (Inactive, red), and reference antimicrobial peptides from curated databases (AMP databases, blue). Reference datasets include CAMP, DRAMP, ADP3, DBAASP, and AMPDB. Descriptor values were normalized to a 0–1 range for comparability across groups.

Case-by-case comparisons further emphasize the importance of residue arrangement. For instance, AI.PBH_19_024 (KKKKRKIKKGKGIVKKKKK) was active, while the nearly identical AI.PBH_19_025 (KKKHEKIKKGDGIVKKKKK) was inactive (**Fig. 4a and Table S1**). Here, three substitutions R→H, K→E, and G→D were sufficient to eliminate antimicrobial potential. A similar pattern emerged in AI.PBH_19_029 (active) versus AI.PBH_19_031 (inactive), where the introduction of cysteine and acidic residues broke the cationic motif. The AI.PBH_25 family provided a particularly illustrative example. Several members (e.g., AI.PBH_25_109, _110, _113, _114, _016, _027, _030) were active, while others with nearly identical backbones (e.g., AI.PBH_25_018, _021) were inactive. The key differentiator was the preservation of an EKKYKKK-rich N-terminal motif in the active variants, whereas the inactive variants introduced acidic substitutions at critical positions within this motif. This demonstrates that not only residue type but also precise positional arrangement governs antimicrobial function.

Beyond charge distribution, further distinctions emerged when considering hydrophobic content and peptide length. The active peptides contained ∼20–25% hydrophobic residues (I, V, L, A, Y, F), compared with ∼15–20% in inactive peptides, indicating that slight hydrophobic enrichment may facilitate membrane insertion (**Fig. 4a and Table S1**). Likewise, the net charge calculated at physiological pH averaged +8 to +12 for the active group versus +4 to +7 for inactive sequences. Active peptides also preserved longer uninterrupted lysine/arginine stretches (5–7 consecutive cationic residues) compared with only 3–4 in inactive ones. These trends suggest that antimicrobial functionality also arises from a finely tuned balance of strong cationic density with localized hydrophobic motifs, rather than from charge content alone.

Interestingly, tyrosine residues emerged as another differentiating feature across peptide families (**Fig. 4a and Table S1**). In the AI.PBH_25 cluster, nearly all active members preserved one or more tyrosine residues embedded within an EKKYKKK-rich motif, suggesting that tyrosine may act as an anchoring residue contributing to amphipathicity and membrane binding. In contrast, the earlier AI.PBH_19 active peptides were entirely tyrosine-free, relying solely on uninterrupted lysine/arginine motifs. Among the inactive set, tyrosine was either absent or positioned within sequences disrupted by acidic substitutions, indicating that the functional role of tyrosine depends strongly on its local sequence context. These observations suggest that while tyrosine is not universally required for antimicrobial activity, its presence within specific cationic motifs may enhance membrane interaction and contribute to peptide efficacy.

Finally, ranking of the active peptides based on their antimicrobial potency revealed further structure–activity relationships. The top-ranked peptides (ranks 1–4) were exclusively from the AI.PBH_25 family, dominated by EKKYKKK-rich motifs with strategically positioned tyrosine residues (**Fig. 4a and Table S1**). These sequences maintained both high cationic density and moderate hydrophobic enrichment. Mid-ranked peptides (ranks 5–8), including AI.PBH_19_024 and AI.PBH_19_029 as well as members of the AI.PBH_39 family, relied on uninterrupted lysine/arginine stretches but lacked the presence of tyrosine motifs, consistent with moderate potency. The lowest-active peptides (ranks 11–13) were also originated from the AI.PBH_19 family and exhibited shorter cationic runs with no or fewer hydrophobic residues, correlating with reduced activity. This gradient underline that while cationic enrichment is a prerequisite for activity, the most potent peptides balance lysine/arginine continuity with tyrosine-mediated amphipathicity and optimized charge–hydrophobic ratios. An additional noteworthy observation is the repetition of tyrosine residues in the most potent peptides. While single tyrosines were occasionally observed in other sequence families, the highest-ranking members of the AI.PBH_25 family consistently contained two tyrosines distributed close to each end of the peptides. This repeated occurrence may provide multiple anchoring points to the bacterial membrane, enhancing amphipathicity and facilitating insertion, likely contributing to their highest activity.

### Silicon exaptation and biophysical basis of emergent antimicrobial functionality

To evaluate whether ML-designed peptides occupy the same biophysical landscape as canonical AMPs, we compared six physicochemical and structural descriptors across active ML peptides (13 peptides), inactive ML peptides (39 peptides), and a reference set of 81,424 AMPs curated from publicly available databases including CAMP, DRAMP, ADP3, DBAASP, and AMPDB (**Fig. 4b**). Database-derived AMPs showed broad distributions reflecting their heterogeneous evolutionary origins. In contrast, ML-designed peptides were restricted to narrower windows, consistent with the imposed design constraint of structural responsiveness. Quantitatively, both active and inactive peptides were far more hydrophilic (mean of about 0.85–0.9 vs. 0.51 in AMPs), exhibited greater polarity (0.73–0.76 vs. 0.25), and were more solvent accessible (0.79–0.83 vs. 0.50) than most reference AMPs (**Fig. 4b**). They also displayed higher pKa values (0.42–0.55 vs. 0.14) and β-sheet propensity (0.7–0.8 vs. 0.55). Bulkiness remained comparable across groups (∼0.61). Within the designed set, active peptides separated from inactive by showing slightly higher pKa and solvent accessibility and slightly lower polarity, suggesting that subtle physicochemical adjustments distinguish functional from non-functional sequences (**Fig. 4b**). We next assessed how many designed peptides fell within the interquartile range of database AMPs for each descriptor. Strikingly, less than 15% of active and inactive peptides exhibited distribution overlaps with AMPs in hydrophilicity, polarity, and solvent accessibility, highlighting that ∼85–90% of ML-designed sequences reside outside the canonical AMP range. By contrast, bulkiness showed >70% overlap, indicating this property is less discriminatory. These results confirm that most ML-designed peptides explore non-canonical sequence space yet still produce antimicrobial activity.

### Selective activity of bacterial membranes and absence of cytotoxic effects on mammalian cells

Because of the striking similarities between biological membranes, since glycerophospholipids are found both in the inner leaflet of bacterial outer membranes and in eukaryotic bilayer membranes, therapeutic use of antimicrobial peptides must target microbial membranes specifically, avoiding harmful interactions with mammalian cells. Therefore, we sought to confirm the cytocompatibility of the lead peptide (AI.PBH_25_016). Human fibroblasts were cultured for 24 h in the presence of peptide and compared with untreated controls and positive control treated with 1% Triton X-100. Fluorescence imaging revealed normal fibroblast morphology, with intact actin cytoskeletons and nuclei indistinguishable from untreated cells (**Fig. 5a**). Consistently, WST-8 assays showed no reduction in metabolic activity following peptide treatment (**Fig. 5b**). Quantitative morphometric analysis further demonstrated that nuclear and cytoplasmic areas were preserved across conditions (**Fig. 5c**), and nuclear shape including roundness, circularity, and aspect ratio remained unchanged (**Fig. 5d**). Together, these results confirm that the peptide is non-cytotoxic to mammalian fibroblasts, supporting its specificity for microbial membranes and its potential suitability for biomedical applications.

**Figure 5.**
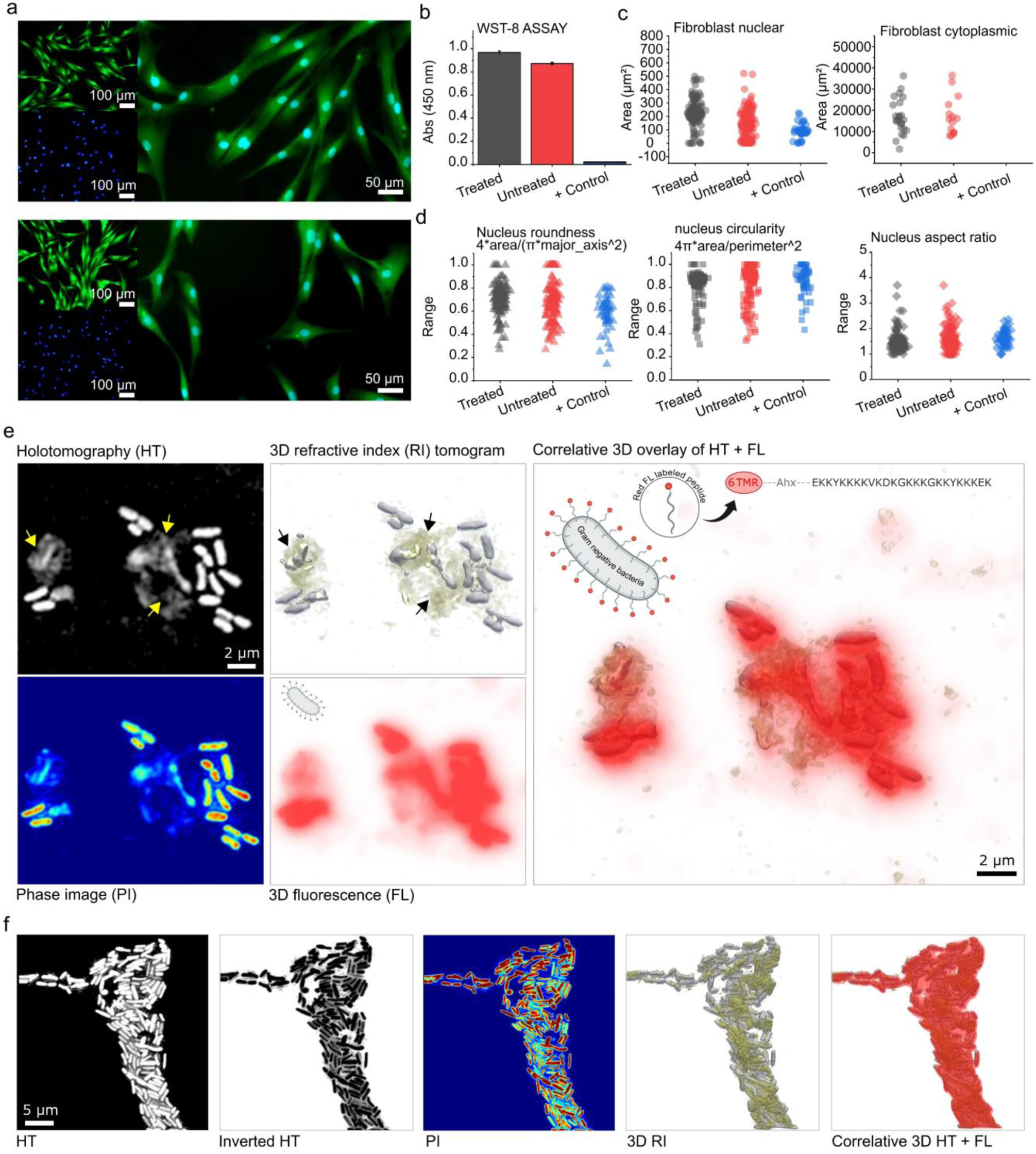
Cytocompatibility of the peptide with fibroblasts and correlative 3D imaging of peptide–bacteria interactions and inhibition. **a)** Representative fluorescence images of human fibroblasts after 24 h exposure to the first ranking peptide AI.PBH_25_01c (“Treated”, top) and untreated (bottom). Actin/cell body is shown in green and nuclei in blue (insets). The cytoplasm of the cell was labeled green, while the nucleus was labeled blue. **(b)** Cell metabolic activity by WST-8 assay (A₄₅₀). Bars show mean and standard deviation across treated, untreated, and +Control conditions. **(c)** Single cell morphometrics of fibroblast nuclear area (left) and cytoplasmic area (right) extracted from fluorescence images. **(d)** Nuclear shape descriptors such as roundness, circularity, and aspect ratio show no adverse changes upon peptide treatment relative to untreated cells. **(e)** Correlative label-free holotomography (HT) and 3D fluorescence (FL) imaging of bacterial cells incubated with lead 6-TMR–labeled (red fluorescence) lead peptide AI.PBH_25_01c. Left: HT amplitude/phase images with yellow arrows marking ruptured bacterial cells and the remaining debris. Middle: 3D refractive-index (RI) tomograms and corresponding 3D FL signal (red) of the labeled peptide. Right: 3D overlay (HT + FL) showing peptide accumulation on/around bacterial cells, consistent with membrane engagement. **(f)** Correlative imaging of cell level bacterial aggregation after incubated with the lead peptides.

### Surface saturation, membrane-targeted rupture and populations collapse

We next investigated how the lead designer peptides specifically interact with and disrupts bacterial membranes. To this end, we performed correlative three-dimensional (3D) HT and fluorescence imaging using a 6-TMR–labeled (red fluorescence) derivative of the top-ranked peptide, AI.PBH_25_016. HT amplitude and phase images revealed ruptured bacterial cells and debris, after peptide exposure (**Fig. 5e, Video S1**). 3D refractive index reconstructions further highlighted structural collapse of bacterial morphology. Corelative 3D fluorescence imaging confirmed strong accumulation of the red labeled peptide at bacterial surfaces and within disrupted cells, consistent with direct membrane binding (**Fig. 5e, Video S2**). The combined imaging data suggests a two-step mechanism, in which peptides first accumulate at the bacterial surface before triggering membrane rupture and collapse.

To our knowledge, this represents one of the first demonstrations of AMP–bacteria interactions captured by correlative 3D holotomography and fluorescence imaging. In addition to direct membrane rupture, correlative imaging occasionally revealed bacterial population aggregation, where adjacent cells appeared to be glued through what we speculate to be blend of ruptured cell debris-peptide mixture (**Fig. 5f, Video S3**). These intercellular connections colocalized with refractive-index changes at contact interfaces, suggesting that peptide assemblies may act as adhesive linkers between bacterial cells with strong red fluorescence signal (**Fig. 5f, Video S4**). Such peptide-mediated adhesion likely reflects residual cationic peptide clusters binding anionic outer-membrane components, effectively gluing neighboring cells and simultaneously disrupting the outer cell membrane. This secondary effect may enhance local peptide concentration and promote cooperative lytic activity, representing a transition from individual-cell disruption to cluster-level aggregation and multicellular collapse within bacterial populations.

### *In Situ* tracking of membrane breakdown and remodeling dynamics with nanoscale resolution

To directly investigate in real time the peptide-induced membrane disruption at nanometer resolution on a fast timescale, we then employed high-speed atomic force microscopy (HS-AFM) on two-dimensional (2D) films reconstructed from extracted *E. coli* outer-membrane lipopolysaccharides (LPS) deposited on mica, as we described in our previous reports.^29^ The high-speed dynamic topography identified four temporally and morphologically distinct phenomena (**Fig. 6a-d**): (i) rapid edge dissolution, (ii) edge reordering and height increase (zone 2), (iii) phase conversion and aggregation, and (iv) height modulation across domains. We identified only about 100-120 seconds into addition of the lead peptide, rapid dissolution initiated preferentially at membrane edges, resulting in progressive loss of surface-bound LPS and the formation of discontinuities in minutes (**Fig. 6a, Video S5,** zone 1). In other regions (**Fig. 6b, Video S6,** zone 2), initially eroded edges partially regrew and exhibited a measurable height increase of about 0.5 nm, suggesting local reorganization and thickening of the membrane surface (Fig. S2). Continued imaging of randomly selected areas after peptide addition also revealed the emergence of dispersed, heterogeneous domains, where previously expanded, uniformly fluid areas seems to transform into compact, high-contrast debris, irregular patches, and elongated aggregates characteristic of solid-like gel phases with sharply defined boundaries (**Fig. 6c, Fig. S3** zone 3-6).

**Figure 6.**
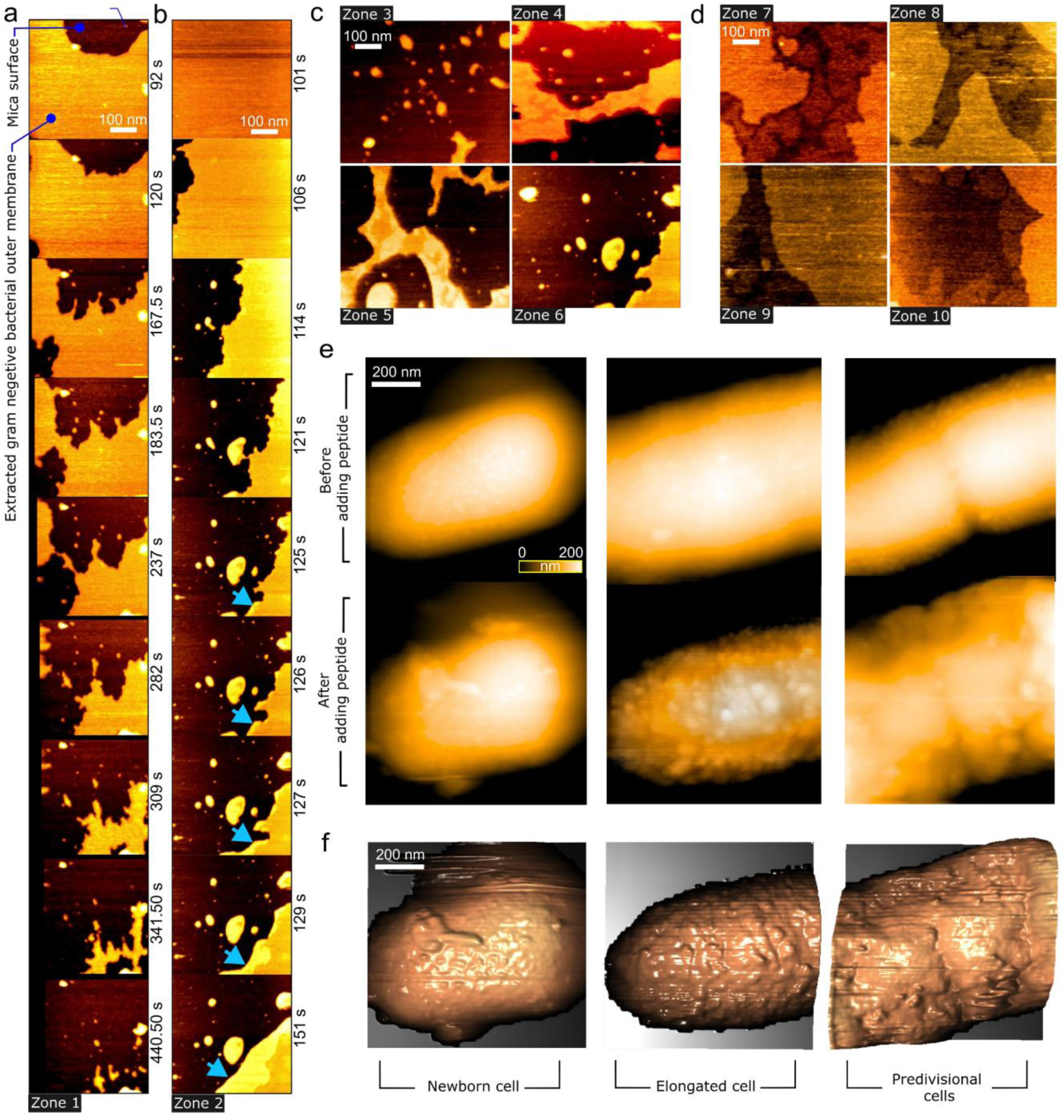
High-speed atomic force microscopy (HS-AFM) reveals AMP-induced blistering and disruption of gram-negative outer membrane and living-cell surface. **a–b)** Time-lapse HS-AFM height images of an extracted gram-negative bacterial outer membrane patch supported on a mica surface following exposure to the top-ranking AMP (Scale bar, 100 nm). **c–d)** Representative HS-AFM height images from additional fields of view (Zones 3–10) (Scale bar, and height profile is 200 nm in all figures). **e)** HS-AFM images of living cells from three morphological subpopulations (newborn, elongated, and predivisional cells) showing identical cells before and after AMP treatment. **f)** 3D reconstruction from the height data corresponding to subpopulations presented in e after peptide treatment (Scale bar is 200 nm in all images).

Further analysis also revealed reproducible vertical height variations ranging from 0.5 to 1 nm contour region across adjacent zones (**Fig. 6d, Fig. S4,** zone 7-10), indicating the coexistence of liquid-crystalline and more ordered β-like membrane. To directly assess the peptide’s impact on the bacterial envelope under physiologically relevant conditions, we then used high-speed AFM on living cells based on our earlier works to visualize real-time cell-surface remodeling upon exposure to the top-ranking AMP (**Fig.6e & f**).^30^ Across three morphological subpopulations newborn, elongated, and predivisional cells, the peptide induced pronounced surface perturbations characterized by the appearance of blister-like protrusions, i.e., localized dome-shaped elevations that interrupt the otherwise continuous surface signal (Fig. e). Higher-resolution topography (Fig. f) resolved these structures as discrete raised domains with altered nanoscale texture relative to the surrounding envelope, consistent with localized delamination and/or compromised envelope integrity. Notably, the extent of blistering increased from newborn to elongated and was most pronounced in predivisional cells, where the surface became progressively more heterogeneous and patchy, suggesting heightened susceptibility to peptide-driven envelope deformation at later stages.

### Molecular dynamics (MD) simulations of Peptide–membrane interactions

To resolve the molecular processes underlying peptide-driven membrane association and remodeling, 1 μs long atomistic MD simulations were performed for top ranking peptide with a mixed POPG/POPE (3:1) bilayer representing a gram-negative outer-membrane mimic. Results showed that the peptides (experimentally relevant concentrations) rapidly adsorbed onto the bilayer within the first ∼0.1 μs and remained predominantly interfacial throughout the 1 μs trajectory (**Fig. 7a**). No apparent transmembrane insertion events were observed throughout the simulation. Instead, peptides primarily accumulated at the lipid–water interface, forming a dynamic interfacial layer that perturbed local lipid packing and increased membrane surface roughness. Mass-density profiles along the bilayer indicate that peptide binding broadens the membrane–water boundary and slightly expands the interfacial region (**Fig. 7b**). Quantitatively, the water–membrane transition width (defined as the distance over which water density decreases from 90% to 10% of bulk) increases from ∼1.18 nm in the peptide-free bilayer to ∼1.61 nm in the presence of peptides (≈37% broader). The lipid headgroup distributions are also modestly broadened on the water-facing side, with outer half-decay widths increasing by ∼12% for POPG and ∼17% for POPE. Peptide density peaks remain localized at ≈ 2.5 nm, with negligible density in the bilayer center.

**Figure 7.**
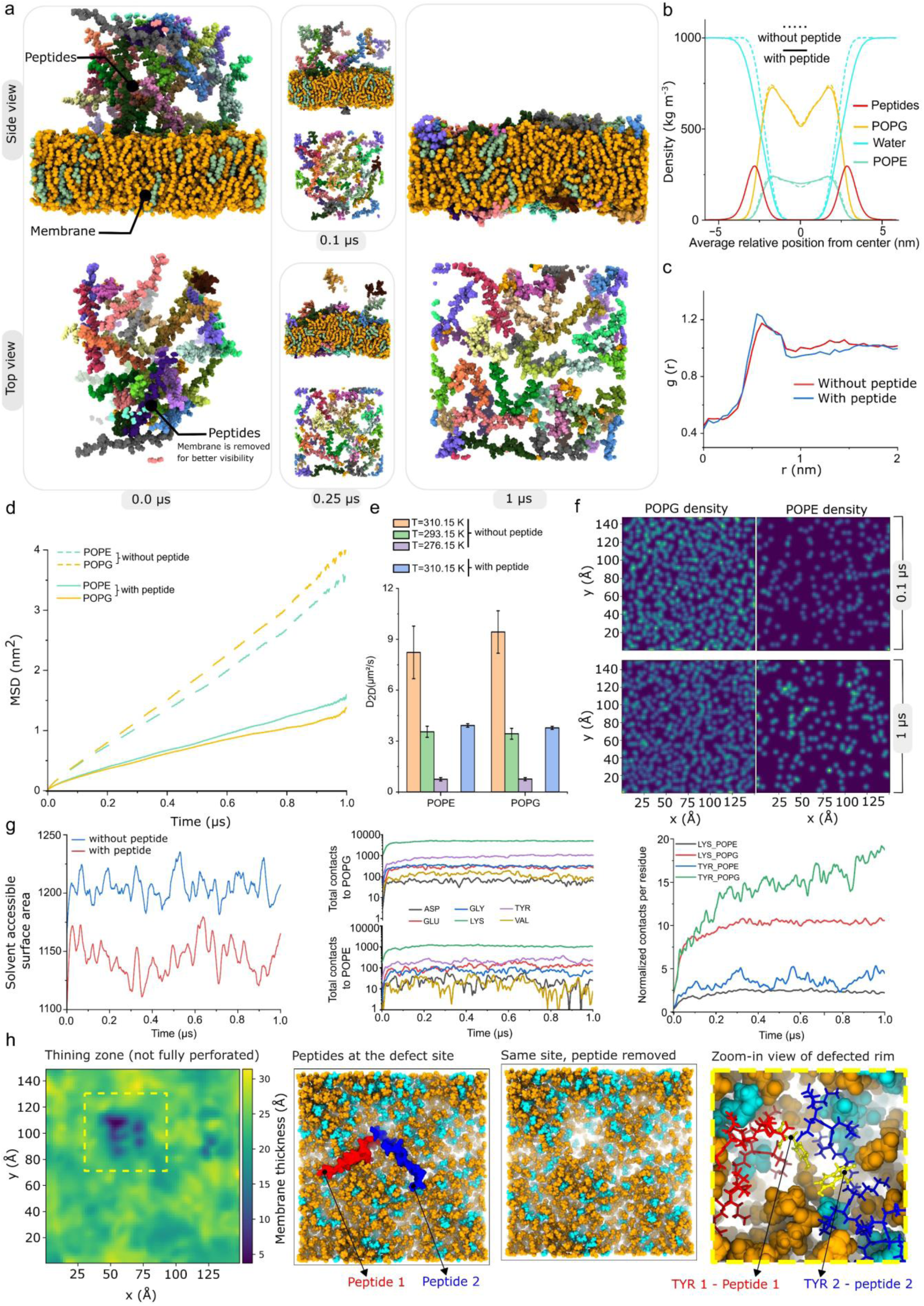
Molecular dynamics (MD) simulations of peptide interactions with outer-membrane model. **a)** Time-resolved MD snapshots showing the evolution of peptides interacting with lipid bilayer. Side- and top-view representations illustrate the progressive adsorption of peptides onto the membrane surface over time. **b)** Density profiles along the bilayer, peptides, and water. Membrane with the peptide is shown with solid line and the results for membrane without peptide are shown as dashed line. **c)** Lateral radial distribution function between POPE headgroups in the presence and absence of peptides. **d)** Mean-square displacement (MSD) of POPG and POPE in presence and absence of peptides. **e)** Lateral diffusion coefficients (D) for POPE and POPG calculated at three temperatures. Peptide binding reduces lipid mobility to a similar extent as lowering the temperature in the peptide-free bilayer, indicating that the peptides impose an ordering/rigidifying effect comparable to membrane cooling. **f)** 2D lateral density maps (Top view) of peptide-driven POPG and POPE redistribution at 0.1 μs and 1 μs. Over time, the bilayer partially develops spatial heterogeneity with local POPE clustering. **g)** Membrane exposure and residue–lipid interaction analysis. Left: solvent-accessible surface area (SASA) of the bilayer with and without peptides, showing reduced solvent exposure upon peptide adsorption, consistent with interfacial reorganization/dehydration observation in panel b. Middle: time evolution of total peptide–lipid contacts to POPG (top) and POPE (bottom), decomposed by residue type, highlighting dominant contributions from cationic and aromatic residues. Right: normalized contacts per residue for Lysine (Lys) and Tyrosine (Tyr) with POPG versus POPE, showing that Lys contacts stabilize early and remain sustained throughout the trajectory, whereas Tyr continues to form new contacts over time. **h)** Defect formation and stabilization. Left: membrane thickness map identifying a local thinning zone not fully perforated (in the center of yellow). Middle: top-view snapshots showing two peptides accumulating at the defect site; removal of peptides at the same site demonstrates the corresponding thinned region. Right: zoom-in of the defect rim highlighting two tyrosine residues from each peptide positioned deep in the middle of the defect zone.

To quantify how adsorption affects membrane dynamics, lipid lateral mobility was assessed via mean-square displacement (MSD) of POPG and POPE (**Fig. 7d**), confirmed quantitatively by increased peak height in lateral distribution function (**Fig. 7c**). In the presence of peptides, MSD strongly suppressed for both lipid species, indicating reduced lateral diffusion. Calculated diffusion coefficients further demonstrated a pronounced rigidification effect (**Fig. 7e**). At 310.15 K, peptide binding reduces D from approximately 8.2 to 3.9 μm² s⁻¹ for POPE (about a 52% decrease) and from 9.5 to 3.8 μm² s⁻¹ for POPG (about a 60% decrease). Notably, this peptide-induced slowdown is comparable to the effect of “cooling” the peptide-free bilayer from 310.15 K to 293.15 K (POPE: ∼8.2 → ∼3.7 μm² s⁻¹; POPG: ∼9.5 → ∼3.5 μm² s⁻¹). This indicates that peptide adsorption imposes an ordering/condensing constraint on lateral lipid motion. That is similar in magnitude to lowering temperature, stronger effect on the anionic POPG component, while non-ionic POPE maintain relatively higher mobility. 2D lipid density maps also provide a qualitative view of lateral organization over the simulation time. The distributions remain broadly mixed for POPE and POPG, with only subtle spatial heterogeneity, arrangements and minor clustering (**Fig. 7f**). We next asked whether surface-bound peptides measurably change how much membrane is exposed to solvent. To quantify peptide-driven changes in interfacial solvent exposure, we computed the membrane solvent-accessible surface area (SASA) which is an integrated readout of interfacial roughness and coverage (**Fig. 7g**, left). SASA drops to 1144.3 ± 18.0 and found to be ∼5% lower than the peptide-free bilayer (1204.7 ± 18.5, Δ=−60.6). Along with density profile (**Fig. 7b**) this indicates that peptides occupy and reorganize the interface, shielding lipid-accessible surface during remodeling to large extent.

To mechanistically link the global membrane response to specific peptide chemistries, we quantified the time evolution of peptide–lipid contacts and decomposed them by residue class and residue-normalized contributions (**Fig. 7g**, middle). The contact landscape strongly gravitated toward two dominant binders, with Lysine providing the largest fraction of peptide– lipid contacts, and tyrosine embedded within the EKKYKKK-rich motif represented the next most prominent contributor. All other residue types contribute comparatively minor contact levels. Importantly, the residue-normalized traces show that Lysine enrichment is not merely a consequence of abundance (**Fig. 7g**, right). Lysine exhibits a sustained binding preference for POPG which nearly stabilized and reached an early plateau, indicative of long-lived headgroup coordination that anchors peptides at the interface. In contrast, tyrosine remains persistently engaged, and progressively maintained contacts with both lipids, but predominantly against POPG. This is consistent with dual functionality of tyrosine, a critical amphiphilic anchor due to its unique dual-nature side chain, which combines an aromatic ring with a polar hydroxyl (-OH) group. This enables dynamic, aromatic-mediated interfacial stabilization, involving interactions such as cation–π interactions with lipid headgroups or adjacent charged residues. The inherent hydrophobicity of the aromatic ring drives its penetration into the hydrophobic acyl-chain region of the lipid bilayer. This persistent interaction allows the tyrosine to continuously sample and stabilize the perturbed membrane interface during membrane remodeling and peptide insertion, which ultimately contributes to membrane disruption, defect or pore formation. Finally, we identified localized thinning defects that remained below full perforation (**Fig. 7h**, right). To better identify these regions throughout our simulation, we calculated a more coarse-grained, density-like measure of bilayer thickness by constructing leaflet surfaces from lipid centre-of-mass (COM) positions.

The representative thickness map illustrated in Figure 7g, is one of several confined thinning events observed over the 1 μs trajectory, where one or more membrane patches locally decreased to ∼10–15 Å compared with a surrounding thickness of ∼20–25 Å (approximately ∼30–40% thinning at the defect core). In the corresponding top-view snapshots, only the two peptides accumulate at this defect site is shown, whereas all the other peptides were removed for better visualization of the phenomena. Removing the peptides at the same location leaves a transiently thinned defected patch, indicating that peptide adsorption can imprint a metastable perturbation in local bilayer architecture (**Fig. 7h**, middle). The zoom-in highlights tyrosine residues in the EKKYKKK-rich motif positioned within the core of thinned zone (**Fig. 7h**, right).

It is noteworthy to mention that our simulations did not progress to clear perforation or large-scale membrane breakup. This likely reflects limitations in trajectory timescale, system size, as well as the slow kinetics of defect growth computationally. Nevertheless, we directly observed strong peptide association with the membrane interface, resulting in increased global lipid rigidity, altered solvation profiles, local thinning events, and partial peptide insertion. All of these points to a defect-like morphology and interfacial reorganization (**Fig. 7**). Collectively, these features are indicative of early-stage lipid destabilization and are qualitatively consistent with the membrane disintegration observed experimentally (**Fig. 6**).

### Hybrid Throttle–Order–Rupture–Collapse (TORC) mechanism

We propose an inhibition mechanism we named as Throttle–Order–Rupture–Collapse (TORC) that reconciles our direct observations from optical microscopy, *in situ* HS-AFM, HS-AFM on reconstructed 2D lipid/LPS films, and atomistic MD simulations. The first step of TORC is that peptides adsorb and enrich at the outer surface. Correlative holotomography-3D fluorescence imaging showed stable interfacial binding (**Fig. 5e**). This is accompanied by suppressed lipid mobility (**Fig. 7d & 7e**) and reduced interfacial solvent accessibility (**Fig. 7b & 7g**) supported by MD, consistent with local condensation/ordering that yields a less hydrated, mechanically stiffer interface rather than immediate detergent-like solubilization or large-area perforation. This ordered state may have two-fold coupled consequences. First, it could impose a transport penalty by restricting diffusion of small solutes and perturbing ion exchange, thereby driving energetic and physiological stress (e.g., reduced PMF/ATP) and diminishing the cell’s capacity to maintain envelope integrity and recover from ongoing membrane remodeling. Second, it progressively loads elastic stress through mechanical mismatch between peptide-rich condensed domains and surrounding more fluid regions, potentially amplified by leaflet asymmetry. Once a critical threshold is reached, the envelope undergoes an out-of-plane instability observed *in situ* as rapid nucleation and proliferation of blister-like domes, followed by coarsening (**Fig. 6e & 6f**). Blister rims and domain boundaries concentrate stress and act as preferential defect sites, consistent with boundary-driven remodeling and edge recession in 2D HS-AFM (**Fig. 6a-c**). As defects proliferate and percolate, catastrophic envelope breakdown yields the collapse phenotype observed by holotomography and MD simulations (**Fig. 5e**, **Fig.6e & 6f, Fig. 7h**). Finally, peptides and released anionic debris can mediate multivalent bridging between cells (**Fig. 5f**), explaining cell–cell aggregation and establishing a positive-feedback loop that locally amplifies both transport limitation and boundary-driven mechanical disruption.

## Conclusion

This work advances a broader view of computational design. The work propose that generative model does not simply produce candidates for one stated objective. It produces a manifold of structured molecular hypotheses. In typical pipelines, only a narrow slice of manifold is retained for the intended task, while outliers or low-ranking targets are discarded as waste or error. Here we propose both the off-objective tail and the successful set are treated as a recoverable resource, because they encode constraints, correlations, and mechanistic pre-existing exaptation learned during training that may be valuable beyond the original goal, where similar physicochemical property profiles matter. Silicon exaptation reframes screening and characterization as an investment that can be reused across applications. Once a model has generated sequences and experiments have produced phenotypes, the resulting output is new information that did not previously exist. Therefore, the question is no longer limited to whether a design meets the original objective, but in addition what the generated space reveals about alternative, emergent uses. At the general level, silicon exaptation may define a transferable workflow. Design for one objective, then interrogate the full output of the model rather than only what passes the primary filter. Identify functional outliers, but also coherent clusters whose shared physicochemical profiles suggest transfer to adjacent applications. Extract minimal, testable rules and feed them back into the next design cycle. This turns generative modeling from single-objective optimization into a discovery engine that harvests value across the entire design space. As model capacity and sampling scale increase, progress will depend not only on choosing better targets, but on making better use of what is already generated. In that sense, computational by-products become starting points for new biology, new materials, new functionalities and use.

## Material and methods

### Computational modeling and sequence generation

Peptide sequences were generated using the updated AIMS2.0 framework, a deep learning– based platform for de novo design of β-hairpin motifs.^2,6,28,31^ The system builds on the earlier AIMS1.0 model and integrates three complementary modules: AIMS-GATHER, AIMS-GENERATE, and AIMS-PROT. In first stage, AIMS-GATHER compiles structural and sequence information from public repositories, identifying homologous folds when possible. If no close relatives are detected, hydrogen-bonding patterns and backbone dihedral angles (Φ and Ψ) are derived using DSSP.^32^ These parameters provide the training basis for the generative and predictive components of the workflow. The second stage, AIMS-GENERATE, is a convolutional neural network (CNN) trained on ∼95,000 protein entries (≤750 residues each). In addition to backbone geometry, the model incorporates physicochemical descriptors, including hydrophobicity, polarity, charge, solvent accessibility, bulkiness, and β-hairpin propensity. By sampling across these feature spaces, the model proposes novel peptide sequences consistent with the targeted secondary structure. Sequence length is constrained by the specified fold, while filters for property thresholds, redundancy, and novelty are applied prior to downstream evaluation. In final stage, AIMS-PROT functions as a structural decoder, reconstructing secondary structures and estimating dihedral angles for candidate sequences. Only sequences that meet accuracy cutoffs advance to molecular dynamics (MD) validation. Each candidate undergoes an initial energy minimization followed by short relaxation simulations to assess conformational stability. Sequences failing to maintain the intended β-hairpin conformation are excluded. Novelty checks against UniProt ensure that retained designs are distinct from existing proteins. All scripts, models, and training datasets associated with the AIMS2.0 workflow are openly available via GitHub.^31^

### Peptide synthesis

All the peptides used in this study including all the original predicted candidates, cysteine mutants and fluorescent labeled listed in **Table S1** were custom synthesized by GenScript (Piscataway, NJ, USA) at about 4 milligrams per sequence using solid-phase peptide synthesis as described previously.^28^ Following TFA-mediated cleavage/deprotection, products were exchanged into the acetate salt and purified; identity and purity were verified by analytical HPLC and mass spectrometry, with ≥ 75% purity. Solubility of the peptide was tested against water, 1X DPBS and 0.1 M PBS.

### Culture media preparation

Mueller-Hinton Broth (MHB) was prepared by dissolving 10.5 g of powdered MHB (Merck, Cat. 70192) in 500 mL of MilliQ water, followed by autoclaving at 121 °C for 15 minutes. Mueller-Hinton Agar (MHA) was prepared by adding 8.5 g of BD Difco™ Granulated Agar (BD Diagnostic Systems) to the broth prior to heating and autoclaving. The agar media was cooled to 50 °C before pouring plates (20 mL/plate) under sterile conditions.

### Bacterial Strain and Cultures

*Escherichia coli* (VTT E-97835) was obtained from the VTT culture collection on nutrient agar and stored at 4 °C. For stock cultures, a single colony was cultured in 5 mL trypticase soy broth (TSB, BD Diagnostic Systems) at 37 °C, 250 rpm for 17 hours. Stocks were stored at −80 °C in 25 % (v/v) sterile glycerol. For working cultures, bacteria from the glycerol stock were streaked onto trypticase soy agar (TSA, BD Diagnostic Systems), incubated at 37 °C for 17 hours, and stored at 4 °C. From that, a single colony was cultured in 5 ml of MHB at 37 °C, 250 rpm for 17 hours. Fresh cell suspensions were diluted in MHB to an optical density at 600 nm (OD600) = 0.1 (∼5 × 10⁵ colony-forming units (CFU)/mL) before experiments. All procedures were performed aseptically.

### Solution-Based Assay

Antibacterial activity of the peptides against E. coli was tested using a solution-based assay. Lyophilized peptides (0.5 mg) from -20 °C were dissolved in 0.2 mL sterile MilliQ water. Different volumes (0.05, 0.025, 0.01, or 0.005 mL) of each peptide solution were added to central wells of 96-well sterile, flat-bottom plates (Falcon Cat. CLS353072), followed by previously described diluted fresh bacterial suspension (OD600 = 0.1) to reach a total volume of 0.1 mL/well. Controls were positioned on each microplate; Positive controls contained ampicillin (100 µg/mL) in MHB, and negative controls contained MHB instead of the peptide solution. MilliQ water was added to the surrounding wells to minimize evaporation. Working volume for each well was 0.1 mL. Plates were sealed using sterile, clear gas-permeable film (Azenta Cat. 4TI-0516/96), then incubated at 37 °C, 300 rpm, for 16 hours in a microplate reader (Varioskan™ LUX, Thermo Scientific™). OD600 was measured at 5-minute intervals.

### Disc Diffusion Assay

The antimicrobial activity was also tested using a disc diffusion assay. From a diluted fresh E. coli suspension (OD600 = 0.1), 0.1 mL was spread onto MHA and allowed to dry for 15 min. Ethanol-sterilized, dried 5 mm filter paper discs (Whatman GF/D 90 mm Cat. 1823090) were placed aseptically on the inoculated agar. Lyophilized peptides (0.5 mg) from -20 °C were dissolved in 0.05 mL sterile MilliQ water and applied onto the blank agar-plated disc. Controls were positioned on a separate plate; Ampicillin solution (400 µg/mL, sterile MilliQ water) served as positive control, and sterile MilliQ water as a negative control. The plates were incubated at 37 °C for 16 hours in an incubator (Innova 44 R, New Brunswick). Antibacterial activity was determined by measuring the zones of inhibition (ZOI) around the disc.

### Cell Lines

Human normal lung fibroblasts (WI-38) were cultured in DMEM high glucose (11965092, Gibco) with Fetal Bovine Serum (FBS, A3840001, Gibco, 10%) and Penicillin/Streptomycin (15070063, Gibco, 1%) at 37°C in a humidified incubator with 5% CO2.

### Cytotoxicity Assay – CCK-8 (WST-8) Assay

Cell viability and proliferation of WI-38 cells following treatment with peptide (Seq. 23_25_016) were evaluated using the CCK-8 (WST-8) colorimetric assay kit (Sigma, 96992). WI-38 cells were seeded in a 96-well plate at a density of 1×10⁴ cells/mL. Experimental conditions included peptide-treated cells, untreated controls, and a positive control group treated with 1% Triton X-100. Each condition was assayed in triplicate.

Upon reaching approximately 70% confluency, peptide was added to the peptide-treated group, while culture medium or 1% Triton X-100 was added to the untreated and positive control groups, respectively. After 24 h, media were removed, cells were washed twice with 1x PBS, and 10% CCK-8 solution was added. The plate was incubated for 3 h at 37 °C in a humidified incubator with 5% CO2, after which absorbance was measured at 450 nm using a microplate reader. Wells containing only CCK-8 solution or culture medium served as blanks. The mean absorbance values of triplicates were calculated, and background values from blanks were subtracted.

### Circular dichroism (CD)

For the CD measurements, the peptides were dissolved in Milli-Q water at a concentration of 0.5 mg/mL, and 300 μL of the peptide solution was placed in a QS quartz cuvette with a path length of 1 mm (Hellma Analytics). The CD spectra was measured using a Chirascan™ CD equipped with a temperature-controlled unit. The CD spectra was measured at 23 °C and prior to the measurement, the sample was allowed to equilibrate inside the chamber for 5 min. The CD spectra was recorded from 190 to 250 nm with a bandwidth of 1 nm, a step size of 1 nm, and a sampling time of 4 s per point. Each CD measurement was repeated three times, and the spectra were averaged before background subtraction (Milli-Q water alone, also measured three times and averaged) and smoothing (factor of 5).

### Fluorescence Imaging

Fluorescence imaging was performed to assess cellular morphology and viability in peptide-treated, untreated, and positive control groups. Experimental conditions were prepared as described above, and treatments were applied for 24 h. Following peptide treatment, cells were labeled with CellTracker™ Green CMFDA (ThermoFisher Scientific, 10644013). Briefly, culture medium was removed, and cells were washed once with 1X PBS prior to incubation with 5 µM cell-tracker solution for 30 min at 37°C in a humidified incubator with 5% CO₂. After incubation, the dye solution was removed, and cells were washed twice with 1X PBS, followed by fixation with 4% paraformaldehyde for 20 min at room temperature. Fixed cells were subsequently counterstained with Hoechst 33342 (ThermoFisher Scientific, H1399) dye for nuclear visualization. Fluorescence images were acquired using a high-NA objective with appropriate filter sets for FITC/GFP (CellTracker™ Green) and DAPI/Hoechst (nuclear counterstain). Images were processed using standard background subtraction, and identical acquisition parameters were applied across all experimental groups.

### Holotomography

Digital holotomography was performed on a Tomocube HT-2H configured as a Mach–Zehnder interferometer with a digital micromirror device (DMD) to scan illumination angles as described previously. ^33^ A coherent 532-nm, 10-mW laser was divided by a 2×2 single-mode fiber coupler into object and reference paths; the specimen arm was imaged with a 60×/1.2 NA water-immersion objective and 175-mm tube lens, together with a matched 60×/1.2 NA water condenser. The two beams were recombined at a beam splitter and the resulting spatially modulated interferograms (holograms) were recorded on a CMOS detector. Volumetric reconstructions from 49 angles used phase-retrieval to obtain amplitude/phase followed by a Rytov-based inversion with the Fourier diffraction theorem to generate 3D refractive-index maps. Theoretical lateral and axial resolutions from the Lauer criterion were 110 nm and 360 μm, respectively. Corelative fluorescence imaging was also performed on the same setup in epi-fluorescence mode using the same objectives and identical conditions. To collect the image excitation from a multi-band LED passed through the appropriate filter cube and dichroic; emitted light was collected through the same objective, filtered for red fluorescence, and recorded on an sCMOS camera, with z-stacks deconvolved and co-registered to the 3D refractive-index tomogram. Samples were prepared in similar was as solution-based assay.

### Molecular dynamics (MD)

Simulations were carried out using GROMACS 2024.2^34,35^ with the CHARMM36^36^ force field (CHARMM-GUI format) and the TIP3P (CHARMM) water model.^37^ The system consisted of 28 identical copies of the peptide embedded in a POPG/POPE (3:1) lipid bilayer, fully solvated and neutralized with 0.15 M NaCl, with no constraints limiting interactions, the peptides could freely contact either the upper or lower leaflet. The CHARMM-GUI Web server (Membrane Builder) was used to generate the membrane.^38–40^ For comparative purposes, an analogous reference system without peptides was also prepared, containing only the lipid bilayer with the same composition and ionic concentration. Each system was subjected to energy minimization, equilibration, and production MD runs. Long-range electrostatics were treated using the Particle Mesh Ewald (PME) method^41^, and van der Waals interactions were modeled using a Lennard–Jones potential with a 1.2 nm cutoff. Bonds involving hydrogens were constrained using the LINCS algorithm^42^, and the geometry of water molecules was maintained using SETTLE.^43^ The simulations were performed with an integration time step of 2 fs for a total of 500 ns for systems without peptide and 1 μs with peptide. The temperature was maintained at 310.15 K, while peptide-free membranes were controlled at 310.15 K, 293.15 K and 276.15 K using the velocity-rescale (Bussi)^44^ thermostat (τₜ = 1.0 ps), and the pressure was kept at 1 bar using the C-rescale barostat in a semi-isotropic coupling scheme (τₚ = 5.0 ps, compressibility of 4.5 × 10⁻⁵ bar⁻¹). The analysis of radial distribution functions (RDF), density distribution, mean squared displacement (MSD), solvent accessible surface area (SASA), as well as number of contacts were performed using built-in Gromacs tools.

Bilayer thickness and lipid lateral organization were calculated from relevant structural snapshots using MDAnalysis^45^, by constructing leaflet-resolved, gridded fields in the membrane plane. POPG and POPE lipids were identified by residue name, and leaflets were assigned based on the sign of the lipid *z*-coordinate relative to the bilayer midplane (defined from the global lipid *z*-distribution in each snapshot), such that lipids above (below) the midplane were classified as upper (lower) leaflet. For COM-based thickness, the mass-weighted center of mass (COM) of each lipid was computed from all atoms, and upper and lower leaflet COM *z*-values were projected onto a regular (*x*, *y*)grid (80 × 80 bins spanning the simulation box) to reconstruct continuous leaflet surfaces. Per-bin leaflet surfaces were estimated from the mean *z* of COMs in that bin, completed for sparse bins by interpolation/nearest-neighbor filling, and smoothed with a 2D Gaussian filter (σ = 1.0, grid units). The COM-based vertical thickness map was then defined as the local surface separation, 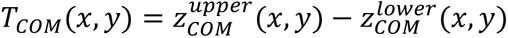.

To quantify POPG/POPE clustering, lipid headgroup phosphorus atoms (P) were used as anchor points for lateral density. For each leaflet and species, P-atom (*x*, *y*)coordinates were binned into a 2D histogram on the same 80 × 80 grid and converted to an areal number density by dividing by bin area; density maps were smoothed with the same Gaussian kernel (σ = 1.0). Total (upper and lower leaflets-combined) densities were obtained by pooling together both leaflets prior to binning. Local composition was expressed as the POPG fraction, *f_POPG_*(*x*, *y*) = *ρ_POPG_*(*x*, *y*)⁄[*ρ_POPG_*(*x*, *y*) + *ρ_POPE_*(*x*, *y*)], computed separately for upper, lower, and total bilayer maps. Identical parameters (lipid definitions, grid resolution, and smoothing) were applied across all snapshots to enable direct spatial correlation between thickness and density/composition patterns.

### High speed atomic force microscopy (HS-AFM)

HS-AFM imaging of supported lipid bilayers was performed using a conventional HS-AFM operated in liquid.^46^ Olympus AC7 cantilevers were used (nominal spring constant 0.2 N/m, resonant frequency ∼800 kHz in liquid, Q factor ∼2). The cantilevers were equipped with electron-beam-deposited (EBD) carbon tips^47^; the same EBD-tip configuration was used for the tip-scan HS-AFM system described below to ensure consistent tip–sample interaction across experiments. All HS-AFM measurements were performed at room temperature.

Supported lipid bilayers were prepared on freshly cleaved mica using polar lipid extract from *E. coli* (Avanti Polar Lipids). Lipids were dissolved in chloroform, dried under N₂ gas, resuspended in 5 mM MgCl₂ to a final lipid concentration of 1.5 mg/mL, and briefly tip-sonicated to form unilamellar lipid vesicles. Subsequently, 2 µL of the vesicle suspension was deposited onto mica, incubated for 5 min, and the surface was thoroughly rinsed to form supported lipid bilayers. HS-AFM imaging was conducted in Milli-Q water. During HS-AFM imaging, peptide solution was injected into the liquid cell. In representative sessions, 0.5–1.0 µL of peptide solution was injected to the 70 µL solution pool, yielding final peptide concentrations typically 7–14 µg/mL. Time-lapse HS-AFM imaging was used to monitor peptide-induced membrane remodeling in real time, including pore formation/expansion and lipid redistribution on the mica surface. HS-AFM movies were processed using an in-house Python-based analysis pipeline (pyNuD).

To analyze in situ surface structure and corresponding mechanical properties, *E. coli* cells exposed to antimicrobial peptides, we utilized high-speed in-line force mapping (HS-iFM)^30^, implemented on a custom-built, tip-scanning HS-AFM integrated with a fluorescence microscope^48^. Olympus AC10 cantilevers equipped with electron-beam deposited carbon tips (tip radius ∼5 nm) were used. Cantilevers were calibrated using the thermal noise method^49^; based on three independent calibrations, the average spring constant was 0.17 ± 0.06 N/m, the resonance frequency 492 ± 91 kHz, and the Q factor 1.65 ± 0.14. *E. coli* strains (NBRC3972/ATCC8739) were acquired from the NBRC and grown in Lysogeny Broth (LB) for 24 h prior to imaging. To immobilize cells, Matsunami NEO cover glass was rinsed with water to remove particulates. A localized observation well was formed by adhering hydrophobic Teflon tape with a 5-mm punched hole to the glass using epoxy. The well surface was coated with poly-L-lysine (PLL; 0.83 g/L, MW 70,000–150,000) for 10 min, followed by incubation of bacterial suspensions on the coated surface for 10 min. Unattached cells were removed by washing with 200 µL of observation buffer. HS-iFM enables simultaneous acquisition of topography and force maps at different spatial resolutions. Topography was recorded at 150 × 150 pixels and force maps at 50 × 50 pixels, with a frame acquisition time of 15 s/frame. Force mapping used a 5 µs loading time and a 300 nm extension, yielding a loading rate of 600 µm/s. Due to the absence of significant adhesion forces, force–indentation curves were fitted using the Hertz model to calculate an apparent elastic modulus. The modulus was averaged over a central region of the cell comprising at least 300 pixels. ^50,51^ All imaging was conducted in 10 mM HEPES, 10 mM NaCl, and 10 mM glucose. HS-AFM movies and force-mapping data were processed using the pyNuD.

## ACKNOWLEDGEMENTS

This work was supported by the Academy of Finland Grant No. 348628, 352900 as well as internal funding from the VTT Technical Research Centre of Finland and Photonics Research and Innovation (PREIN) flagship. P. B. and G.W. acknowledge the financial support of the statutory research fund of ICSC PAS. T.U. acknowledges support from MEXT Promotion of Development of a Joint Usage/ Research System Project: Coalition of Universities for Research Excellence Program (CURE) Grant Number JPMXP1323015482 and JSPS KAKENHI Grant Number 24K01309. The solid-state NMR studies were supported by the EU project Fragment-Screen (grant agreement ID: 101094131). B. F.-Y. acknowledges the postdoctoral funding from the Sustainable Biomedical and Toxicological Research (SUSBIO) - Profiling action on soft materials. The authors wish to acknowledge CSC – IT Center for Science, Finland, as well as Poland’s high-performance computing infrastructure PLGrid (HPC Centers: ACK Cyfronet AGH) grant no. PLG/2025/018882, for providing computational resources.

## SUPPORTING INFORMATION

Additional supporting information can be found online in the Supporting Information section at the end of this article.

## Supporting information

**Table S1.**
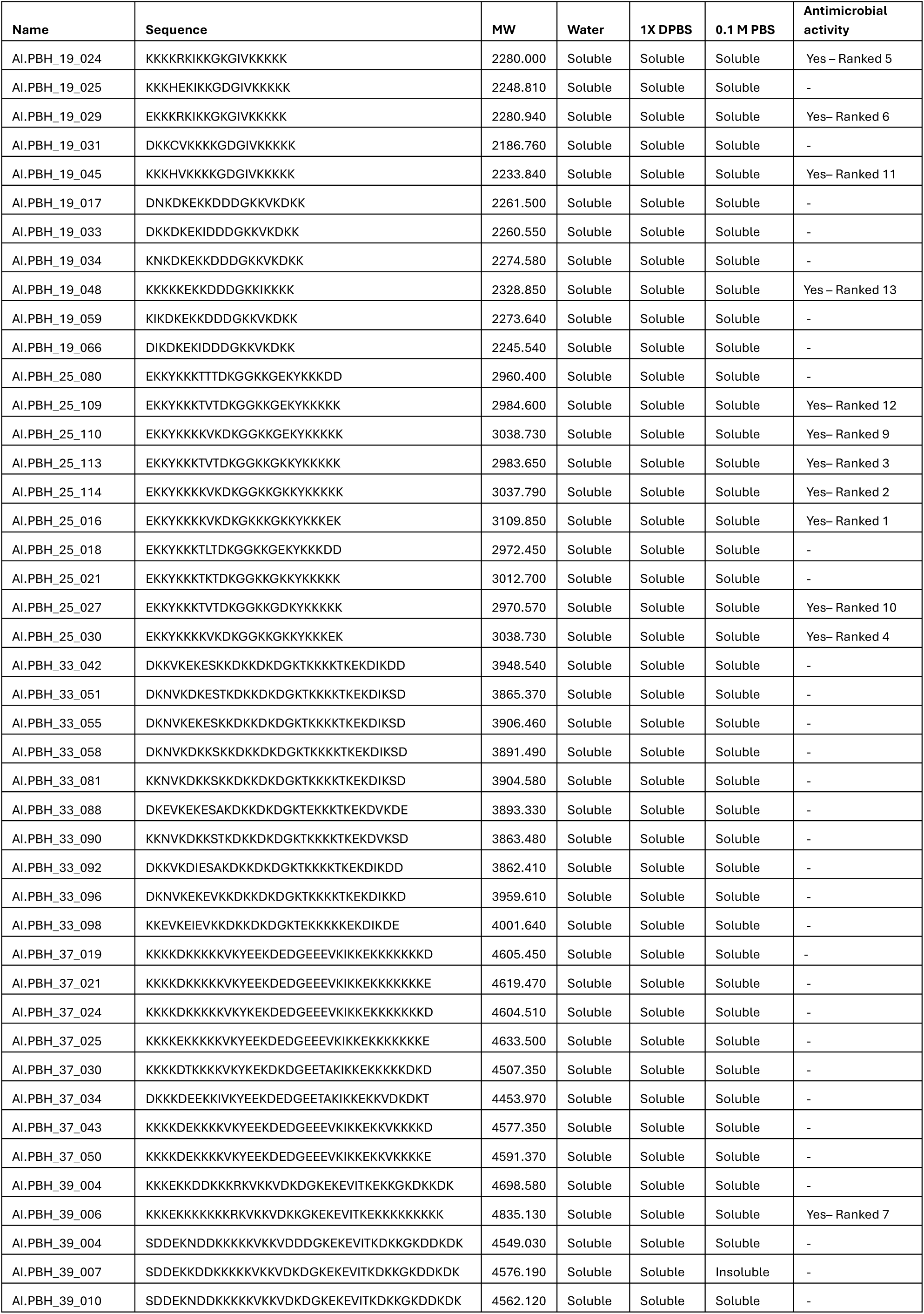
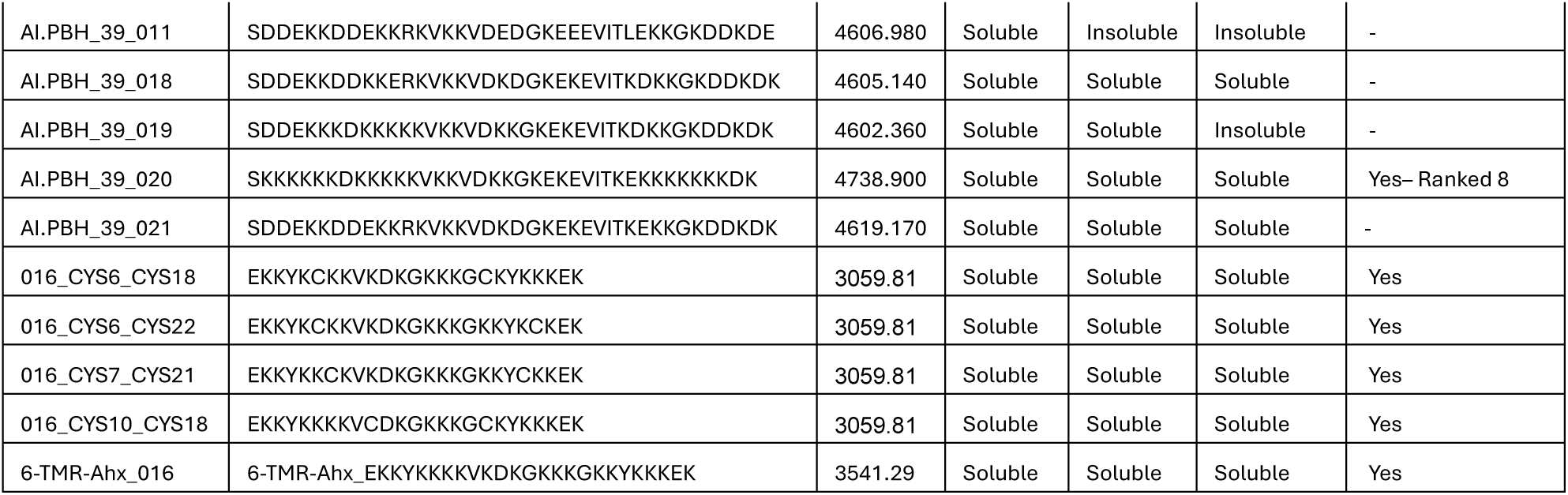
AIMS 2.0 predicted *de novo* designed peptides, their sequence information, solubility in different solution and antimicrobial properties.

**Table S2.** Simulated systems comprising peptides embedded in a POPG/POPE (3:1) lipid bilayer and a reference system without peptides.

|  |  |  |  |  |  |  |  |  |
| --- | --- | --- | --- | --- | --- | --- | --- | --- |
| ID | nPOPE | nPOPG | nprotein | nwater | nNa | nCl | Temperature | final box size [nm3] |
| Membrane & peptide | 179 | 537 | 28 | 115238 | 457 | 312 | 310,15 K | 15.22 x 115.22 x 19.16 |
| Membrane | 184 | 552 | 0 | 29642 | 629 | 77 | 310,15 K | 15.38 x 15.38 x 7.52 |

**Supplementary Figure 1.**
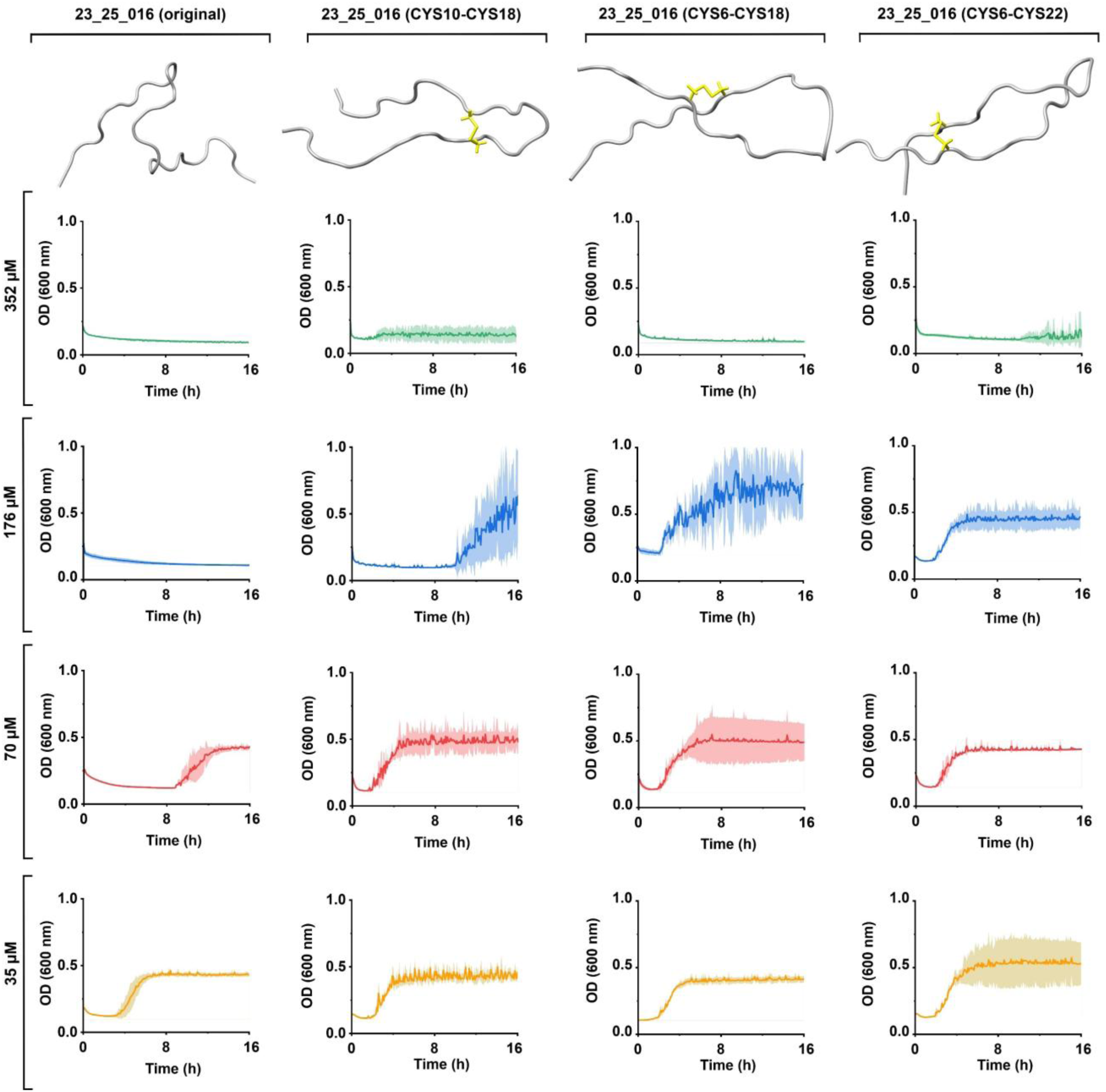
Growth kinetics in the presence of peptide 23_25_016 and disulfide-locked variants. Representative backbone conformations of the parent peptide (23_25_016, original) and three engineered disulfide connectivities (Cys10–Cys18, Cys6– Cys18, and Cys6–Cys22) are shown at the top; disulfide bonds are highlighted in yellow. Bacterial growth was monitored as optical density at 600 nm (OD 600) over 16 h in the presence of each peptide at the indicated concentrations (352, 176, 70, and 35 µM; rows). Solid lines show the mean OD 600 and shaded bands indicate variability across replicates (*n* = 3 in all cases).

**Supplementary Figure 2.**
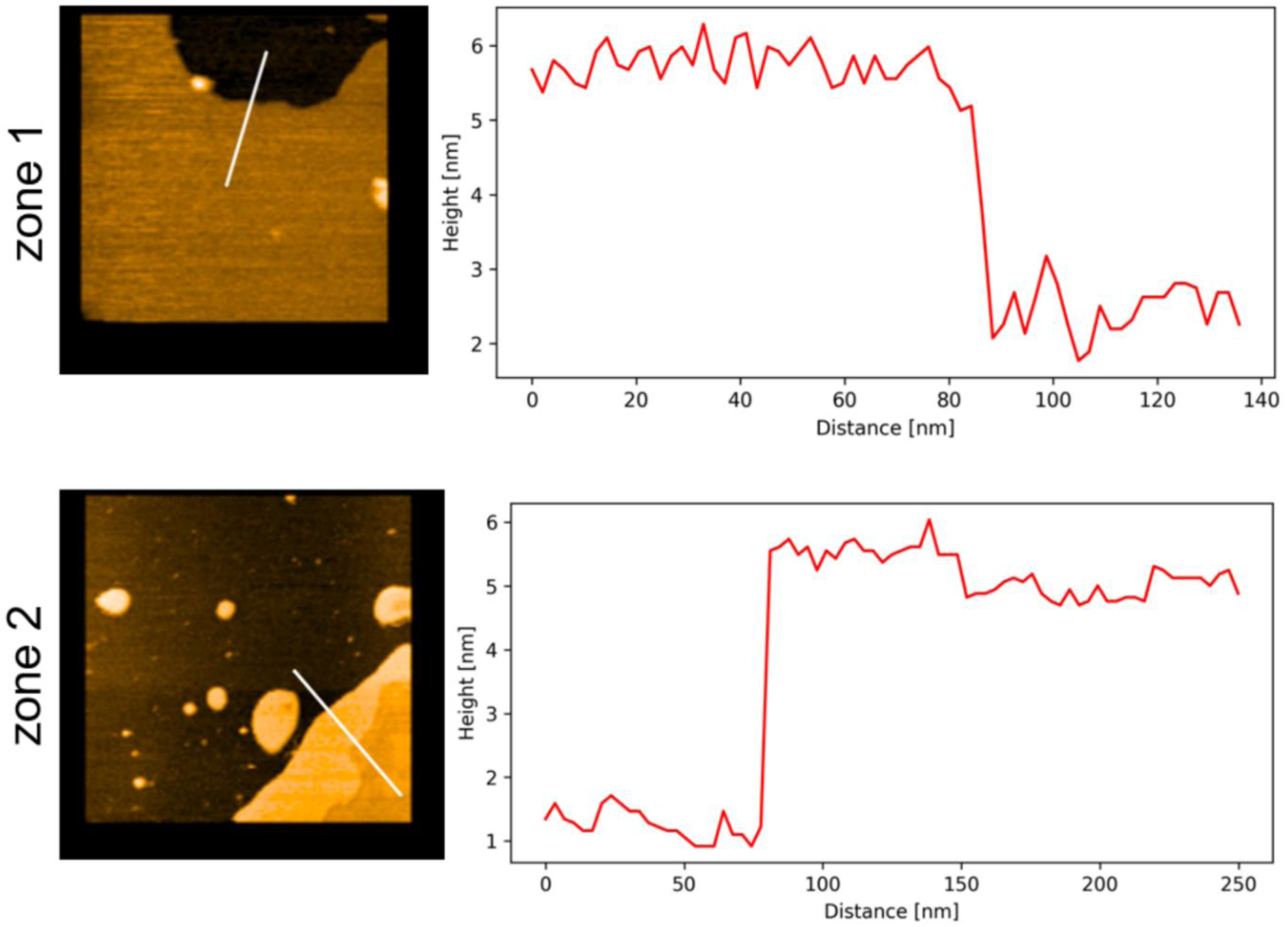
HS-AFM topography and representative height profiles of Gram-negative lipid patches on mica (Zones 1–2). HS-AFM height images (left) show Gram-negative extracted lipids spread on a mica substrate, where darker regions correspond to exposed mica and brighter regions correspond to lipid-covered areas and lipid-associated features. The white line indicates the transect used to extract the corresponding height profile (right; red trace), plotted as height (nm) versus lateral distance (nm).

**Supplementary Figure 3.**
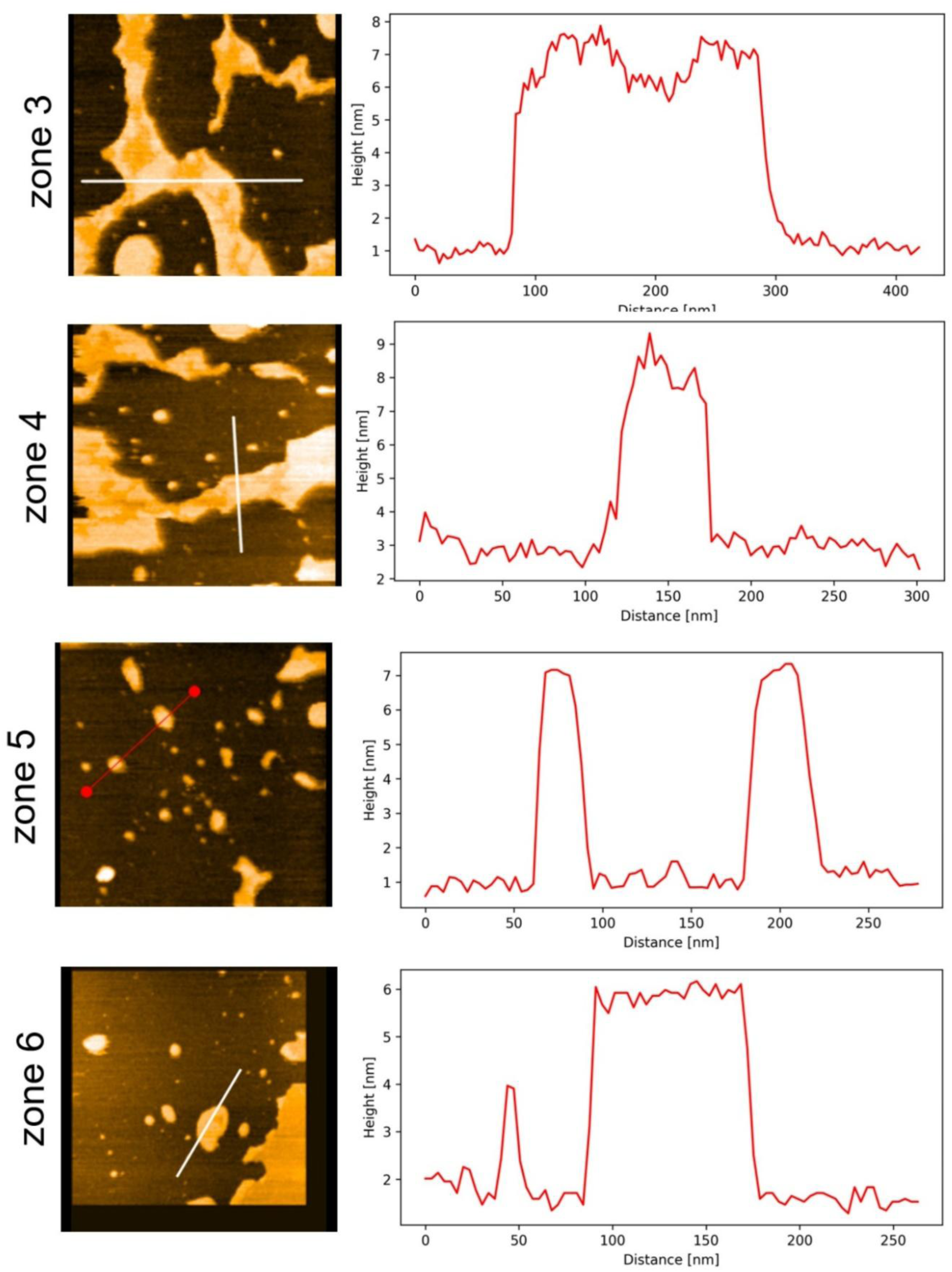
HS-AFM topography and representative height profiles of Gram-negative lipid patches on mica (Zones 3–6). HS-AFM height images (left) show Gram-negative extracted lipids spread on a mica substrate, where darker regions correspond to exposed mica and brighter regions correspond to lipid-covered areas and lipid-associated features. The white line indicates the transect used to extract the corresponding height profile (right; red trace), plotted as height (nm) versus lateral distance (nm).

**Supplementary Figure 4.**
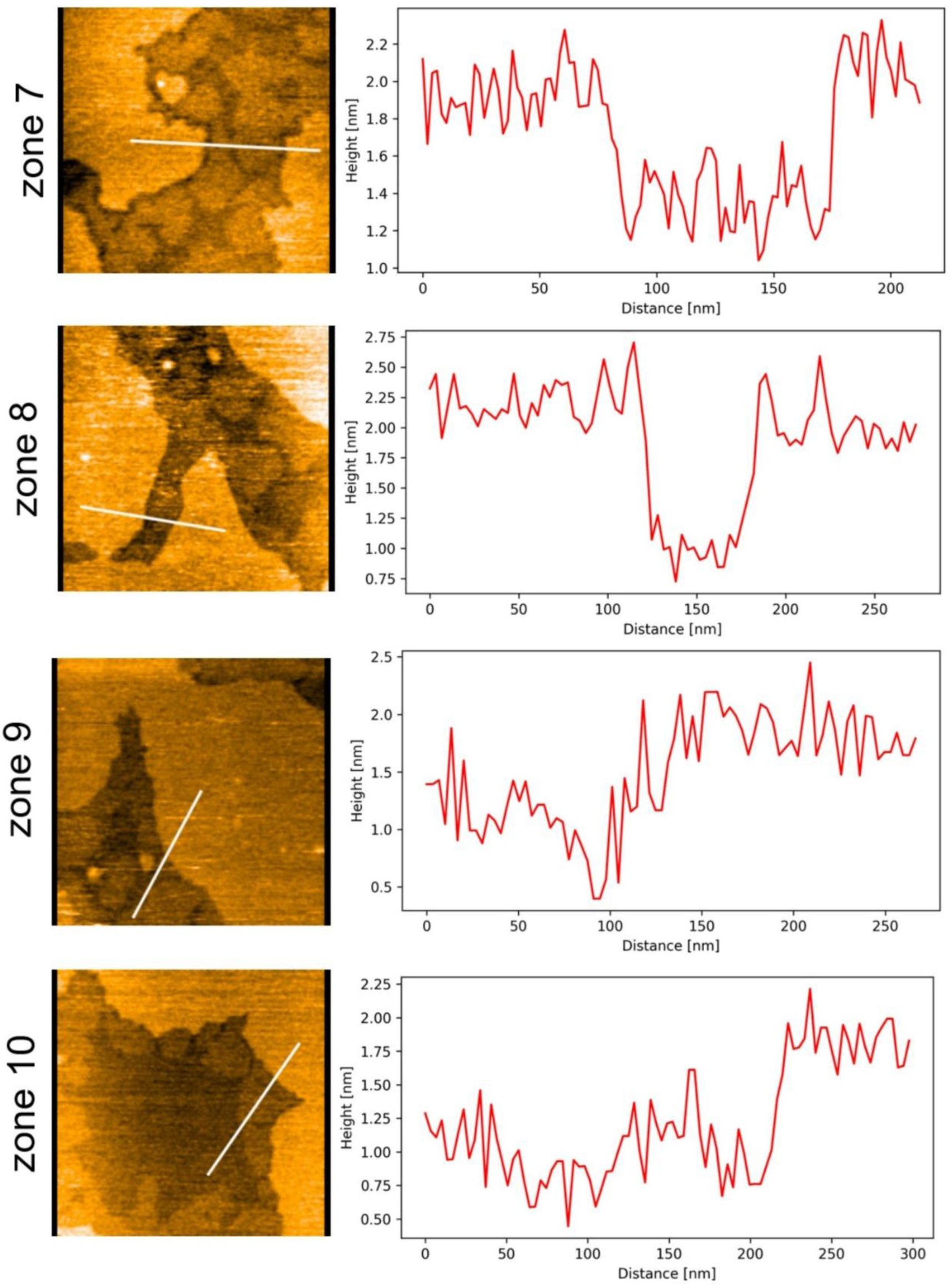
HS-AFM topography and representative height profiles of Gram-negative lipid patches on mica (Zones 7–10). HS-AFM height images (left) show Gram-negative extracted lipids spread on a mica substrate, where darker regions correspond to exposed mica and brighter regions correspond to lipid-covered areas and lipid-associated features. The white line indicates the transect used to extract the corresponding height profile (right; red trace), plotted as height (nm) versus lateral distance (nm).

